# Bacteriophage self-counting in the presence of viral replication

**DOI:** 10.1101/2021.02.24.432718

**Authors:** Seth Coleman, Tianyou Yao, Thu Vu Phuc Nguyen, Ido Golding, Oleg Igoshin

## Abstract

When host cells are in low abundance, temperate bacteriophages opt for dormant (lysogenic) infection. Phage lambda implements this strategy by increasing the frequency of lysogeny at higher multiplicity of infection (MOI). However, it remains unclear how the phage reliably counts infecting viral genomes even as their intracellular number increases due to replication. By combining theoretical modeling with single-cell measurements of viral copy number and gene expression, we find that, instead of hindering lambda’s decision, replication facilitates it. In a nonreplicating mutant, viral gene expression simply scales with MOI rather than diverging into lytic (virulent) and lysogenic trajectories. A similar pattern is followed during early infection by wildtype phage. However, later in the infection, the modulation of viral replication by the decision genes amplifies the initially modest gene expression differences into divergent trajectories. Replication thus ensures the optimal decision—lysis upon single-phage infection, lysogeny at higher MOI.

## INTRODUCTION

Following genome entry into the host cell, temperate bacteriophages must choose between two developmental pathways (Ofir and Sorek, 2018). In the default, lytic pathway, rapid viral replication typically culminates in the death of the host cell (lysis) and release of viral progeny. By contrast, in the lysogenic pathway, phages suppress their virulent functions and enter a dormant prophage state (Ofir and Sorek, 2018). To decide on the infected cell’s fate, temperate phages assess the environmental abundance of potential hosts (Golding et al., 2021; Ofir and Sorek, 2018). If susceptible host cells are scarce, then producing hundreds of new phages via the lytic pathway would be futile, and, instead, lysogeny should be chosen. To evaluate the relative abundance of viruses and cells, phages use diverse methods. Some achieve this by measuring the number of simultaneously coinfecting phages (multiplicity of infection, MOI) and increasing the frequency of lysogeny at higher MOI (Bourret and Fox, 1988; Levine, 1957). Other bacteriophages harness cell-cell communication to assess the frequency of virus-host encounters in their vicinity (Erez et al., 2017; Silpe and Bassler, 2019). Notwithstanding the mechanism by which the measurement is performed, a regulatory circuit encoded by the virus must interpret a biological signal reflecting the relative abundance of viruses and host cells and use it to bias a decision between the two possible outcomes of infection.

Phage lambda, a temperate bacteriophage that infects *Escherichia coli*, has long served as the paradigm for viral self-counting (Arkin et al., 1998; Golding, 2011; Kourilsky, 1973; Weitz et al., 2008). Direct measurements, both in bulk (Kourilsky, 1973; Lieb, 1953) and in single cells (Zeng et al., 2010), demonstrated that a higher number of coinfecting phages increases the probability of lysogeny. Decades of experimental interrogation have resulted in a comprehensive genetic understanding of the virus and the identification of key players involved in the lambda postinfection decision (Casjens and Hendrix, 2015; Court et al., 2007; Dodd et al., 2005). However, despite this detailed molecular knowledge of the underlying circuitry, our system-level understanding of how MOI drives the infection outcome is far from complete (Joh and Weitz, 2011; Kobiler et al., 2005; Zeng et al., 2010). In contrast to the two-gene “switch” governing lysogenic maintenance (Ptashne, 2004), the network driving the lysis/lysogeny decision comprises multiple genes, regulating each other through diverse molecular interactions (Casjens and Hendrix, 2015). The common theoretical view of the decision is that this genetic network is biased by MOI towards either of two attractors, one corresponding to lytic onset, another to lysogeny (Arkin et al., 1998; Avlund et al., 2009; Cortes et al., 2017; Joh and Weitz, 2011; Kobiler et al., 2005; McAdams and Shapiro, 1995; Shao et al., 2018; Weitz et al., 2008). However, the way this takes place varies between models. Further challenging our ability to decipher the circuit’s function, and seemingly inconsistent with the two-attractors picture, is the fact that, while the eventual gene expression patterns in lysis and lysogeny clearly differ, the initial gene expression cascade following infection appears indistinguishable for both pathways (Oppenheim et al., 2005).

Complicating phages’ task of measuring MOI—and our attempts to decipher how they do it—is the fact that viral copy number is rapidly increasing inside the infected cell (**Figure 1**). Phage replication begins within minutes of genome entry (Shao et al., 2018) and coincides with the expression of early genes in the decision circuit (Oppenheim et al., 2005)(see **Figure 2** below). In other words, the initial MOI, which the viral circuitry presumably attempts to measure (Gandon, 2016; Sinha et al., 2017), is soon obfuscated by the presence of additional phage genomes in the cell. Elucidating how lambda succeeds in distinguishing the initial genome number from the instantaneous number present in the cell has remained a challenge partly due to experimental limitations. Within a population, single-cell MOI is broadly distributed (Kourilsky, 1973; Zeng et al., 2010)(**Figure S1**), necessitating measurements at the level of the individually infected cell. However, simultaneous measurement of viral copy number and the expression of phage genes has previously not been possible at single-cell resolution.

**Figure 1.**
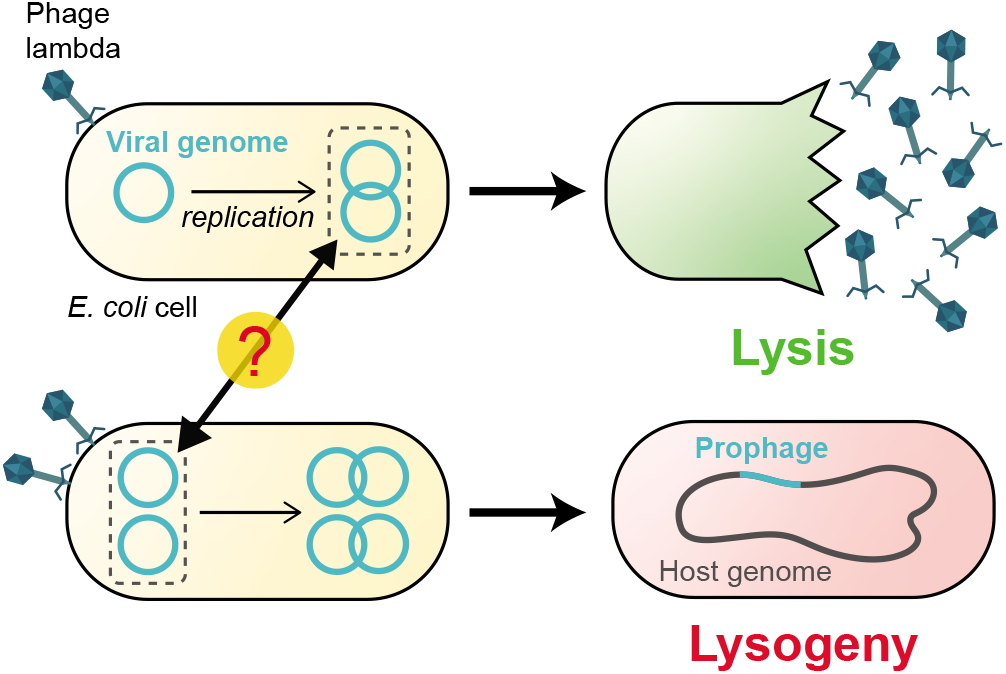
The lambda decision circuit measures the multiplicity of infection (MOI) even as viral copy number is changing. A higher multiplicity of infection (MOI) increases the propensity to lysogenize. Here, infection by a single lambda phage (top) results in lysis, whereas coinfection by two phages (bottom) leads to lysogeny. In choosing cell fate, the infecting phage must respond to the initial number of viral genomes in the cell but ignore the subsequent increase in number due to viral replication.

**Figure 2.**
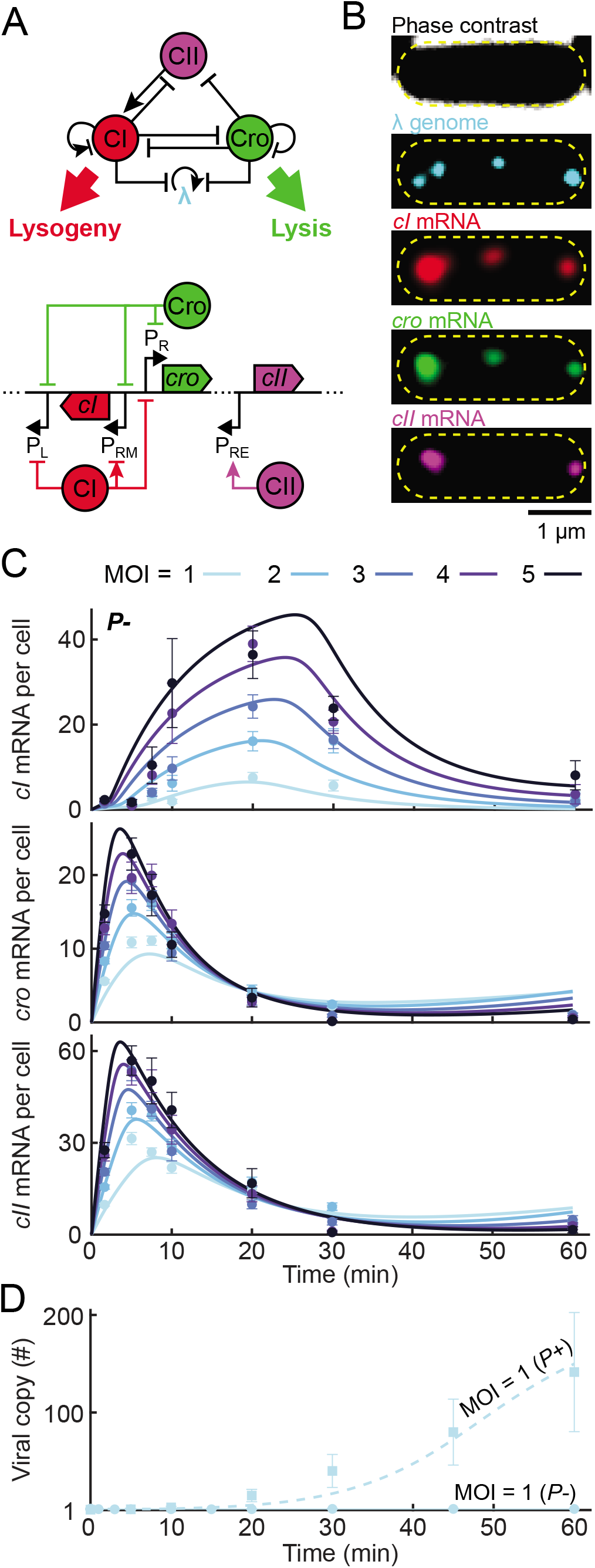
A simplified model of the decision network captures the kinetics of mRNA and viral copy number following infection. (A) Top, the three-gene circuit at the heart of the lysis/lysogeny decision. Bottom, the corresponding segment of the lambda genome. Upon viral entry, P_R_ expresses both *cro* and (following a leaky terminator) *cII*. CII then activates *cI* expression from P_RE_. CI and Cro repress P_R_ and P_L_, as well as phage replication. In a lysogen, CI regulates its own expression from P_RM_. (B) Images of a single *E. coli* cell at 10 minutes following infection by λ *cI*857 *Pam*80 *P1parS*. Phage genomes are labeled using ParB-*parS*, and the mRNA for *cI*, *cro*, and *cII* using smFISH. Yellow dashed line indicates the cell boundary. (C) The numbers of *cI*, *cro*, and *cII* mRNA per cell, at different times following infection by λ *cI*857 *Pam*80 *P1parS*, at MOI = 1–5. Markers and error bars indicate experimental mean ± SEM per sample (see **Table S6** for detailed sample sizes). Solid lines indicate model fit. (D) Viral copy number, measured using qPCR, following infection at MOI = 1 by *P*+ (λ *cI*857 *Sam7)* and *P*- (λ *cI*857 *Pam*80 *P1parS)* phages. Markers and error bars indicate experimental mean ± standard deviation due to qPCR calibration uncertainty. Lines indicate model prediction. See **Experimental Methods** and **Theoretical Methods** for detailed experimental procedures, image and data analysis, and modeling.

Here, we combine single-molecule detection of mRNA and phage genomes during infection with mathematical modeling of network dynamics, to identify how lambda measures the number of coinfecting phages. To circumvent the complication of a time-varying genome number, we first examined infection by a replication-deficient lambda strain. At various times after infection, we measured, in individual cells, the viral copy number (which, in this case, equals the MOI) and mRNA levels of key lambda genes—*cI, cro*, and *cII*. To our surprise, we found no divergence of the mRNA trajectories between low and high MOI, indicative of a transition between the lytic and lysogenic attractors. Instead, gene expression simply scaled with viral dosage. This led us to hypothesize that viral replication is required for obtaining an MOI-dependent lysis/lysogeny decision. To test this hypothesis, we constructed a data-calibrated model for the decision network that included the coupling of phage replication to gene expression. Our model revealed that, indeed, viral replication is inextricably coupled to the lysogeny decision. Early in infection, during a time window set by the dynamics of CII, a short-lived activator, expression of the lysogenic repressor CI scales with MOI, similarly to what we observed in the absence of replication. However, subsequent replication—and its modulation by the decision genes—drive a sharp divergence of cell fates as a function of MOI. Specifically, the initial accumulation of CI at MOI > 1 leads to repression of both *cro* expression and viral replication, enabling the lysogenic choice. In contrast, at MOI = 1, accumulated CI is insufficient to repress *cro* expression and replication. Consequently, *Cro* production from a rapidly increasing number of *cro* gene copies activates the lytic pathway. We thus find that, rather than hindering lambda’s decision by obscuring the initial MOI, viral replication ensures the appropriate choice of lysis upon infection by a single phage and lysogeny upon coinfection.

## RESULTS

### In the absence of viral replication, gene expression does not diverge into lytic and lysogenic trajectories

To characterize the behavior of the lambda decision circuit, we sought to measure gene expression kinetics during infection across a range of single-cell MOI values. To decouple the gene-regulatory aspects from the effects of time-varying dosage, we first followed the approach of Kourilsky (1973) and Arkin et al. (1998) and examined infection by a replication-deficient mutant (*Pam*80, henceforth denoted *P*-)(Kourilsky, 1973). We focused on the expression of three lambda genes at the heart of the decision circuit—*cI, cro*, and *cII* (Avlund et al., 2009; Shao et al., 2018; Weitz et al., 2008)(**Figure 2A**). Cro, transcribed from P_R_, is a repressor that inhibits transcription of multiple lambda genes (including itself) from the two early promoters, P_R_ and P_L_ (Court et al., 2007). Cro is required for successful lysis, to prevent the accumulation of CI and the overproduction of lambda proteins deleterious to later development (Court et al., 2007). Transcription of *cro* can also be used as a proxy for the presence of Q, a critical lytic gene produced from the same polycistronic transcript (Oppenheim et al., 2005). CI, too, inhibits transcription from P_R_ and P_L_, in addition to regulating its own expression from PRM, and is required for establishing and maintaining lysogeny (Oppenheim et al., 2005). CII is a short-lived protein that activates early transcription of *cI* from P_RE_, a critical event for the establishment of lysogeny (Court et al., 2007). Both CI and Cro also suppress viral replication by inhibiting expression of the lambda replication proteins O and P (Oppenheim et al., 2005) and by repressing transcription from P_R_, required for early replication (Casjens and Hendrix, 2015).

To measure the MOI dependence of expression dynamics in this three-gene subnetwork, we combined single-molecule quantification of mRNA and phage genomes in individual cells (Wang et al., 2019)(**Figure 2B**). Following infection by a replication-deficient phage (*cI*857 *Pam*80 *P1 parS*; see **Experimental Methods** for strain construction and experimental protocols), samples were taken at different time points and chemically fixed. The lambda genomes present in each cell were detected and counted using the ParB-*parS* system (Tal et al., 2014; Youngren et al., 2014)(**Experimental Methods** and **Figure S2**). In the same cells, mRNA copy numbers for *cI*, *cro*, and *cII* were measured using single-molecule fluorescence *in situ* hybridization (smFISH) (Skinner et al., 2013; Wang et al., 2019). The cells were then grouped based on the measured single-cell MOI, and the averaged mRNA level for each gene, time, and MOI, was calculated (**Figure 2C**).

All three genes exhibited a transient pulse of expression, with mRNA numbers first rising, then decaying (**Figure 2C**). The main difference between genes was in the timing of the expression peak, with *cro* and *cII* reaching their highest level approximately ~10 minutes after the entry of viral genomes, and *cI* peaking later, at about ~20 minutes. Biological replicates yielded consistent results (**Figure S3**). The observed dynamics were consistent with our current understanding of the gene expression cascade following infection: upon viral entry, *cro* and *cII* are transcribed from P_R_ (see **Figure 2A**) and this promoter is later repressed by Cro (Oppenheim et al., 2005). *cI* transcription requires activation of the P_RE_ promoter and is hence delayed until enough CII protein, driving this activation, has accumulated (Oppenheim et al., 2005).

Previous models of the lysis/lysogeny decision, both in the presence (Avlund et al., 2009; Cortes et al., 2017) and absence (Joh and Weitz, 2011; McAdams and Shapiro, 1995; Weitz et al., 2008) of viral replication, predicted the existence of distinct patterns of gene expression, identifiable as the two possible infection outcomes. In light of this prevailing picture, we were surprised to observe no clear divergence of mRNA trajectories between low and high MOI, reflecting a transition from lysis to lysogeny. Instead, we found that for each of the genes, a simple scaling by a factor MOI^ε^ (suggested previously by Joh and Weitz, 2011), with ε ≈ 1 for *cI* and ε ≈ 0.5 for *cro* and *cII*, yielded a near collapse of the different MOI-gated trajectories to a single curve (**Figure S4**). Numerically integrating the mRNA numbers to estimate the instantaneous concentrations of Cro and CI likewise did not reveal a divergence of trajectories between low and high MOI (**Figure S5**), indicating that the lack of divergence is not merely an artifact of examining the short-lived mRNA species.

### Modeling network dynamics reveals that viral replication is required for a lysis-to-lysogeny transition

The absence of a clear MOI-driven lysis-to-lysogeny transition following infection by a nonreplicating phage led us to hypothesize that viral replication is required for such a transition to take place. To explore this hypothesis, we constructed a deterministic mathematical model that describes the regulatory interactions between *cI*, *cro*, and *cII*, as well as the coupling between gene expression and viral replication (**Figure 2A** and **Theoretical Methods**). Building on previous theoretical efforts (Avlund et al., 2009; Shao et al., 2018; Weitz et al., 2008), our model captures, phenomenologically, both direct interactions between the three genes (e.g., the activation of *cI* transcription from P_RE_ by CII and the repression of *cro* transcription from P_R_ by CI) and indirect ones, mediated by players that are not modeled explicitly (e.g., CIII-mediated suppression of CII degradation—see Kobiler et al., 2007). As its output, the model produces the population-averaged temporal dynamics of the viral copy number and mRNA and protein concentrations, for a given initial MOI. To estimate the parameters governing gene regulation in the network, we fitted the model to the experimental mRNA kinetics during *P*- infection, by minimizing the least-squares error using particle swarm optimization (Kennedy and Eberhart, 1995)(see **Theoretical Methods**). This procedure yielded a good agreement between the model and experiments (**Figure 2C** and **Figure S6**).

To relate the gene expression dynamics computed by our model with the infection outcome, we assumed that cell fate is determined by whether Cro or CI concentration in the cell reaches a threshold value (Joh and Weitz, 2011). The lytic threshold reflects Cro’s role in repressing *cI* transcription from P_RM_ and (indirectly, by repressing *cII* transcription from P_R_) from P_RE_. The Cro threshold also reflects the requirement that Q (encoded by the same transcript as Cro) reaches sufficient level to enable readthrough of the late lytic genes transcribed from P_R’_ (Kobiler et al., 2005). The lysogenic threshold, on the other hand, corresponds to CI levels sufficient to turn off the P_L_ and P_R_ promoters, thereby repressing the expression of lytic genes and phage replication (Oppenheim et al., 2005).

Using our model to analyze the network dynamics following *P*- infection, we found that, consistent with what we inferred from the mRNA data above, the predicted shape of protein trajectories in the CI-Cro plane (**Figure 3A**) was largely unchanged with MOI, with both species merely increasing in concentration with the viral copy number. While a range of CI thresholds can be defined, which ensure lysogeny above some critical MOI, no Cro threshold can result in lytic decision below that critical MOI value (**Theoretical Methods**). Instead, the inferred trajectories suggest that the lytic threshold is never reached during *P*- infection, and that, at low MOI, neither lysis nor lysogeny is selected. This interpretation is consistent with the known absence of lysis following infection by *P*- phages (Brooks, 1965; Shao et al., 2018), but suggests that this failure reflects the state of the decision circuit, rather than merely a failure to execute the chosen lytic pathway, which was the implicit assumption in previous works (Arkin et al., 1998; Joh and Weitz, 2011; McAdams and Shapiro, 1995).

**Figure 3.**
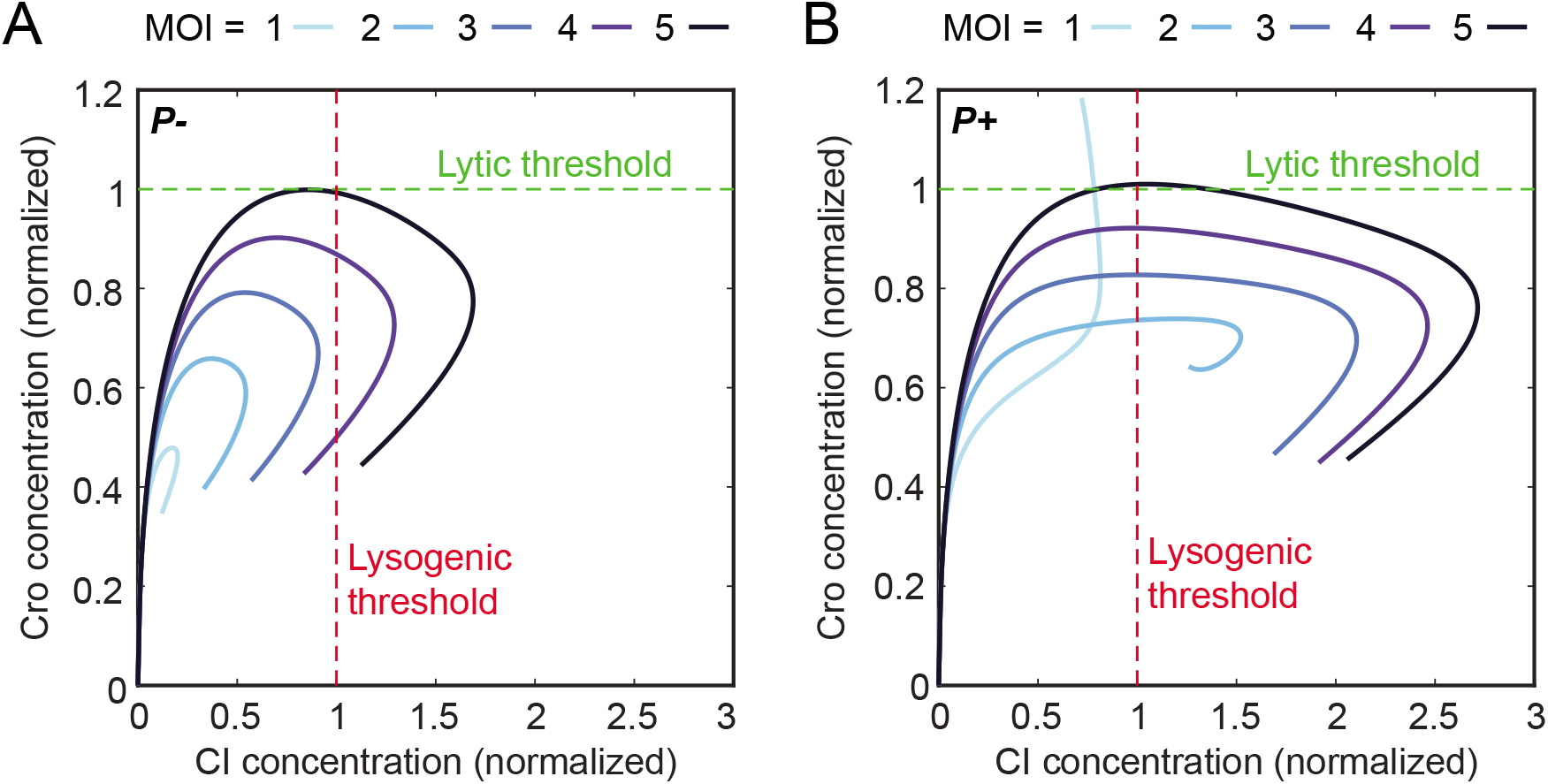
Phage replication is required for an MOI-driven lysis-to-lysogeny transition. (A) Model-predicted trajectories, in the plane of Cro and CI concentrations, during the first 60 minutes following infection by *P*- (nonreplicating) phage at varying MOI. Protein concentrations were normalized by the lytic and lysogenic thresholds. (B) Same as panel A, for the case of infection by a replicating (*P*+) phage. See **Theoretical Methods** for detailed information.

We next sought to evaluate what effect viral replication would have on the system’s behavior. To calibrate the model parameters pertaining to lambda replication and its regulation, we used qPCR measurements of genome number kinetics from low-MOI infection by a replicating (*P*+) phage (**Figure 2D**), as well as published data for *cI* and *cII* expression following infection with *P*+ phages, *cI-, cro*-, and *cro-P*-mutants (Shao et al., 2018)(**Figures S7, S8**). The resulting model allowed us to calculate the CI-Cro trajectories following infection by a replicating phage at various MOIs. In contrast to what we observed for the nonreplicating phage, these trajectories show a clear divergence between MOI = 1 and higher MOIs, and, in particular, support a transition from lysis (Cro threshold crossing) to lysogeny (CI threshold crossing) with increasing MOI (**Figure 3B**). Moreover, a single choice of thresholds is simultaneously consistent with the experimental phenotypes of both *P*- and *P*+ phages in terms of the MOI value at which the transition to lysogeny occurs (≈ 3–4 and 2 for *P*- and *P*+ respectively; see **Theoretical Methods** and **Figure 6** below)(Kourilsky, 1973).

**Figure 6.**
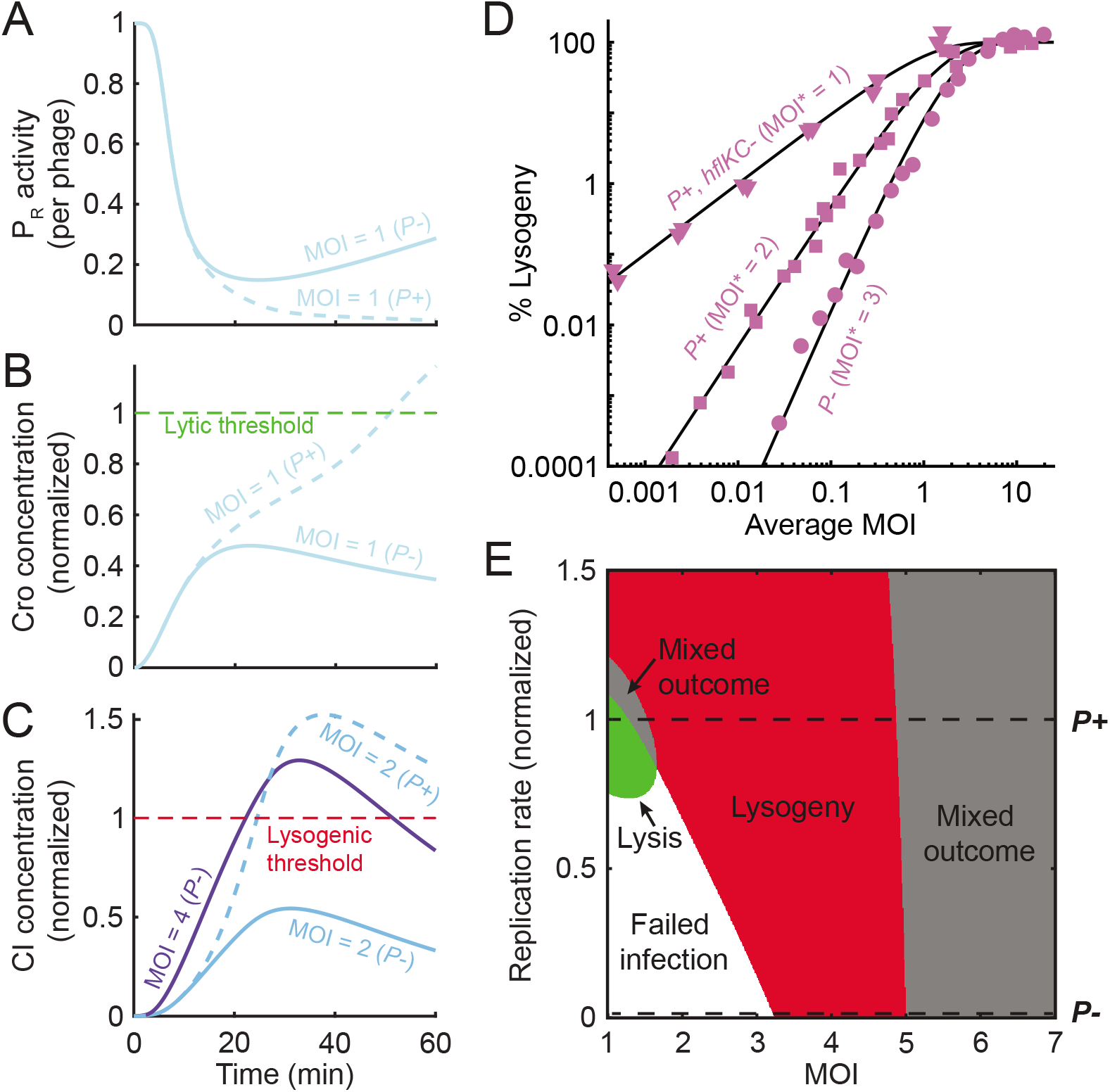
Replication is required for the lytic outcome and lowers the MOI required for lysogeny. (A) Model-predicted P_R_ activity per phage during *P*- (solid blue line) and *P*+ (dashed blue line) infection at MOI = 1. (B) Cellular Cro concentration, normalized by the lytic threshold, for the cases modeled in panel A. (C) Cellular CI concentration, normalized by the lysogenic threshold, during *P*- infection at MOI = 2 and 4 (solid light and dark blue lines, respectively), and during *P*+ infection at MOI = 2 (dashed line). (D) The fraction of cells undergoing lysogeny as a function of average MOI, during bulk infection with *P*- (circles), *P*+ (squares), and phages with prolonged CII lifetime (*P*+ phages infecting *hflKC*- hosts; triangles). The experimental data was fitted to a model (black lines) where virus-cell encounters follow Poisson statistics, and infection at MOI ≥ MOI* results in lysogeny (Kourilsky, 1973). The *hflKC*- strains are either Δ*hflK* or Δ*hflC* (see **Table S1**). (E) Predicted infection outcome as a function of MOI and viral replication rate (normalized by the fitted replication rate for *P*+ phage). See **Experimental Methods** and **Theoretical Methods** for detailed experimental procedures, data analysis, and modeling.

### CII activation of P_RE_ defines a time window for the network’s response to MOI

Having successfully recapitulated the decision phenotype, we next sought to understand how lambda reliably responds to the initial MOI even in the presence of viral replication. Since establishing lysogeny requires reaching a critical CI concentration, we focused on the response of *cI* expression to MOI. Two lambda promoters drive *cI* transcription, P_RE_ (activated by CII) and P_RM_ (autoregulated by CI) (Oppenheim et al., 2005). P_RM_ is solely responsible for CI production in a lysogen (Oppenheim et al., 2005), but whether it plays a role during the initial decision has remained unresolved (Michalowski and Little, 2005). Our model indicates that the MOI-driven increase in CI is caused by transcription from P_RE_, and that removing *cI* autoregulation does not eliminate that response (**Figure S9**). This finding is consistent with reports that a wide range of mutations in P_RM_ permit the establishment of lysogeny (Michalowski et al., 2004; Schubert et al., 2007), whereas mutating P_RE_ or CII prevents it (Oppenheim et al., 2005). Thus, to elucidate CI’s response to MOI, we focused on characterizing the CII-activated expression of *cI* during infection.

Our model indicates that CII-activated P_RE_ expression occurs in a single pulse, taking place within the first ~30 minutes of infection (**Figure 4A**). The amplitude (per phage) and duration of this P_RE_ activation pulse depend only weakly on MOI (**Figure 4A** and **Figure S10**). These predictions are supported by direct measurement of nascent *cI* mRNA level at individual phage genomes (**Figure S11**). They are also consistent with our findings above that, in non-replicating phages, cellular *cII* numbers are dosage compensated whereas *cI* level scales linearly with MOI (**Figure S4**).

**Figure 4.**
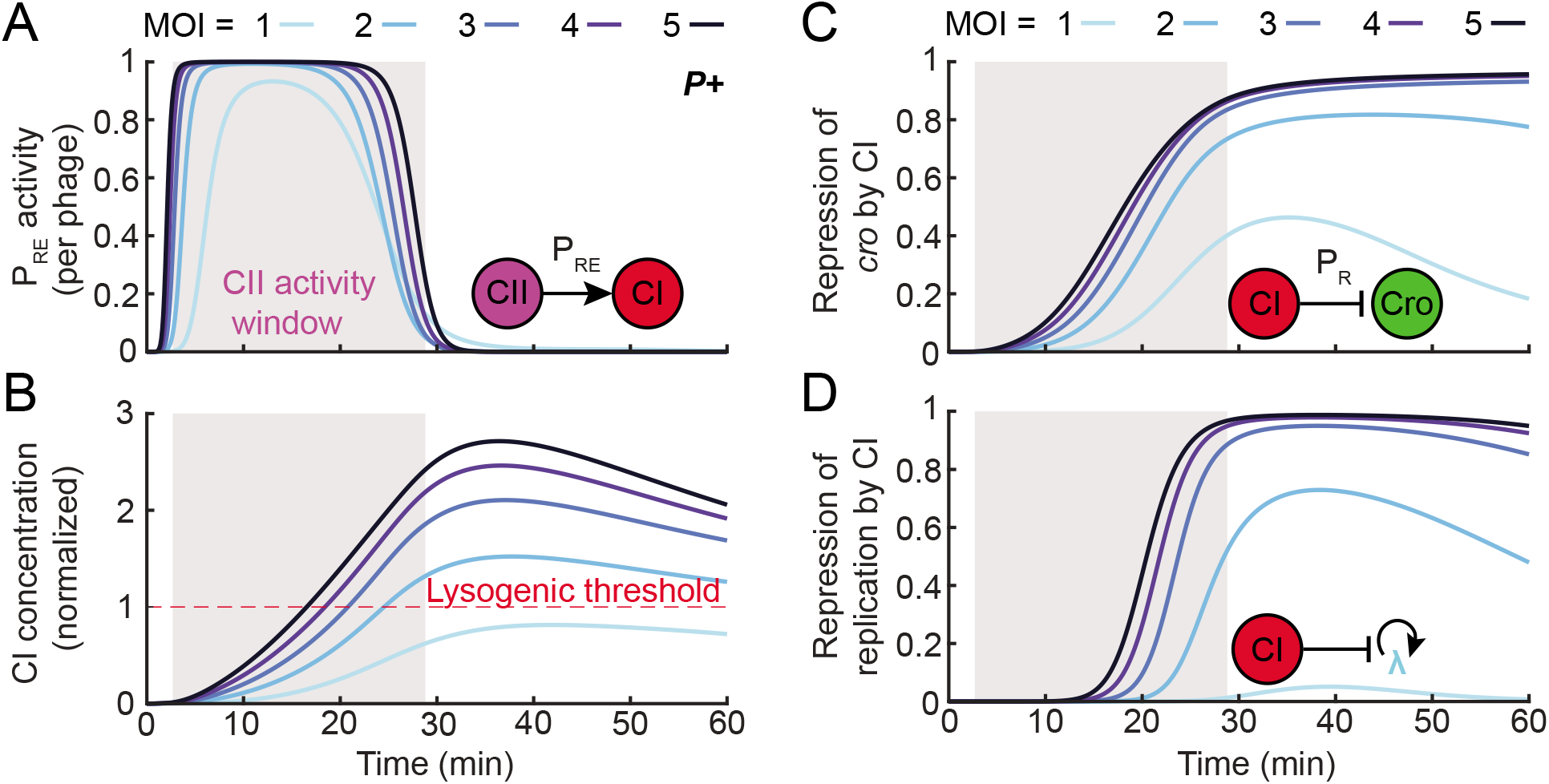
CII activation of P_RE_ defines a time window for the network’s response to MOI. (A) Model-predicted P_RE_ promoter activity (per phage) following infection by *P*+ phage at MOI = 1–5. Gray shading indicates the MOI-averaged CII activity window, defined as the period during which P_RE_ activity is greater than 10% of its maximum. (B) Cellular CI concentration (normalized by the threshold concentration for lysogeny, dashed red line) following infection by *P*+ phage at MOI = 1–5. (C) The strength of *cro* repression by CI, calculated as the magnitude of the CI repression term in the P_R_ transcription rate, following infection by *P*+ phage at MOI = 1–5. (D) The strength of repression of replication by CI, calculated as the magnitude of the CI repression term in the viral replication rate, following infection by *P*+ phage at MOI = 1–5. See **Theoretical Methods** for detailed information.

The MOI independence of *cI* expression from individual viral copies during the P_RE_ activation window provides a necessary element for the lysogeny decision, by guaranteeing that cellular CI concentration increases with MOI. Specifically, the maximum CI concentration at MOI = 2 is approximately 2-fold higher than for MOI = 1, and this fold-change in CI concentration is first reached within the P_RE_ window (**Figure 4B**). Higher MOI further increases CI concentration within the same time window (**Figure 4B**). The end result is that, at MOI ≥ 2, CI concentration reaches the critical level sufficient to repress both *cro* expression (**Figure 4C**) and viral replication (**Figure 4D**), leading to lysogeny. This repression is established during the P_RE_ activation window and persists throughout infection (**Figure 4C & D**).

### Changes in viral copy number outside the CII activity window do not alter the decision

The findings above reveal how CI levels—and the propensity to lysogenize—increase with MOI. However, a reliable MOI-based decision requires also that viral replication inside the cell will not obfuscate the initial response to MOI. We reasoned that, for this to hold, the system should become insensitive to changes in viral dosage once the window for CII activation of P_RE_ is closed. To test this hypothesis, we followed the approach of Cortes et al. (2017) and used the model to examine what happens in the case of delayed infection. Specifically, we modeled an infection by a single phage, followed by a second single-phage infection at time τ_d_ later (**Figure 5**). We found that, indeed, if the second infection takes place after the end of the P_RE_ activity window, τ_pre_, the outcome is lysis, and the CI and Cro trajectories are indistinguishable from those at MOI = 1 (**Figure 5A**). If, in contrast, the delayed infection occurs early enough within the P_RE_ activity window, infection results in lysogeny, with CI and Cro trajectories similar to those for a synchronized infection at MOI = 2 (**Figure 5A**). Infecting with higher numbers of late-arriving viruses lengthens the time window where the late phages can affect the decision, but not beyond τ_pre_, (**Figure S12**). Thus, the outcome is insensitive to changes in viral copy number that take place outside the time windows determined by CII activity.

**Figure 5.**
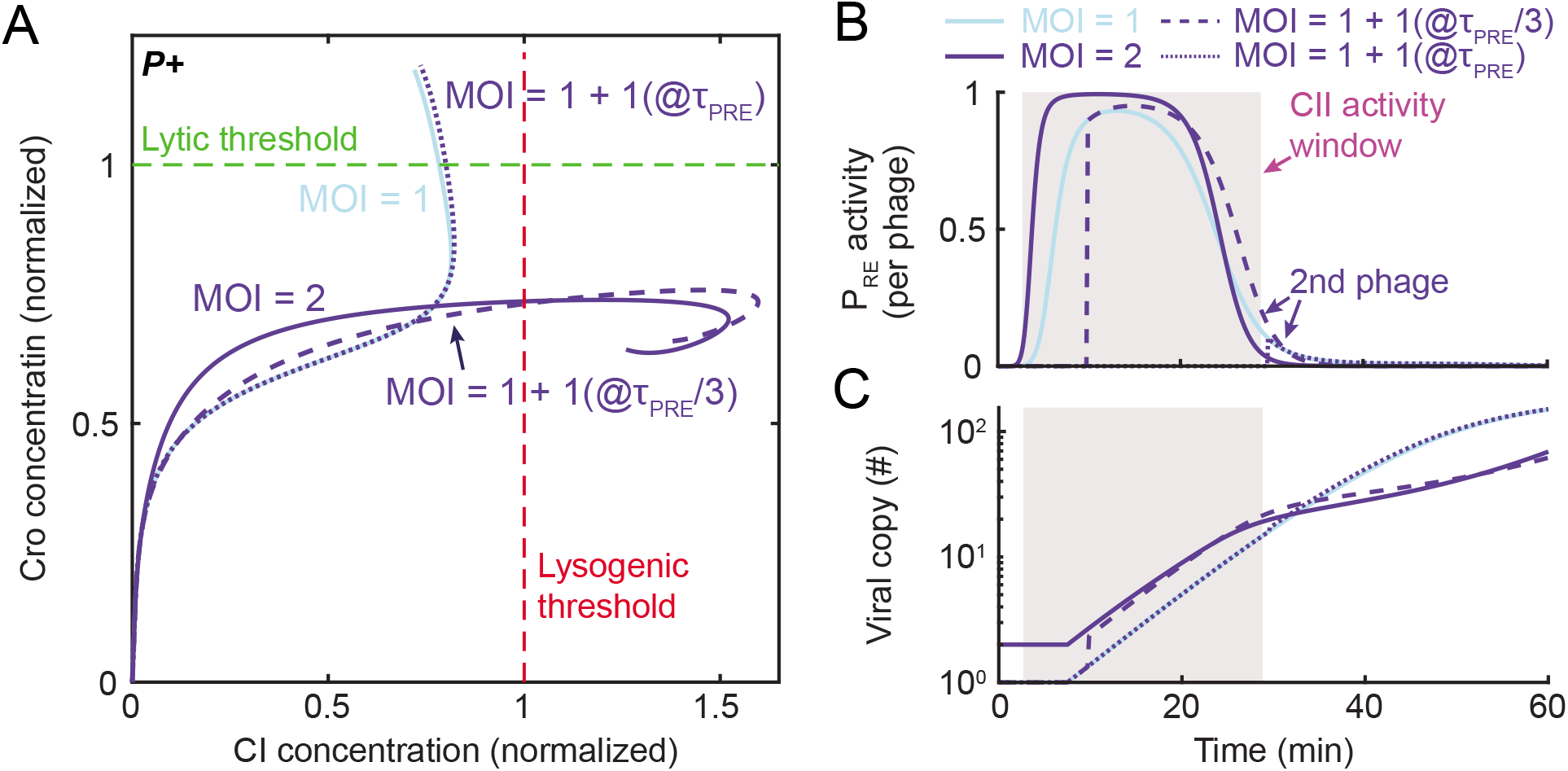
Changes in viral copy number outside the CII activity window do not alter the decision. (A) Model-predicted system trajectories, in the plane of Cro and CI concentration, during the first 60 minutes following infection by *P*+ phages, at 4 different scenarios: Infection by a single phage (solid line, light blue), infection by a single phage followed by a second phage at time τ_PRE_/3 (with τ_PRE_ the end of the CII activity window; dashed line, dark blue) and τ_PRE_ (dotted line, dark blue), and simultaneous infection by two phages (solid line, dark blue). (B) P_RE_ activity from the second arriving phages, for the cases modeled in panel A. The shaded grey region indicates the MOI-averaged CII activity window estimated for synchronized infections. (C) Viral copy number over time, for the cases modeled in panel A. See **Theoretical Methods** for detailed information.

Why does delayed infection result in a diminished response? Our model indicates reduced P_RE_ expression from the second virus, since that virus is not present for the entire duration of the CII activity pulse (**Figure 5B**). This results in lower cellular accumulation of CI during that time window, as compared to simultaneous coinfection (**Figure 5A**). Furthermore, the delay impacts not only the late-infecting virus itself but also all viruses produced subsequently through viral replication (**Figure 5C**). We note that, through the combination of Cro repression of P_R_ and active CII degradation (Oppenheim et al., 2005), the decision network constrains CII activity to a single time window. Subsequent changes in viral copy number after the initial infection cannot overcome this constraint. These features of the system’s response to delayed infection can also explain why rampant viral replication during infection at MOI = 1 does not cause a switch to lysogeny: While replication generates >100 additional genome copies, viruses produced outside the CII activity window are unable to express *cI* from P_RE_ (**Figure S13**). The opportunity to ‘flip’ the decision switch has already been lost.

### Phage replication enables the lytic choice and lowers the MOI required for lysogeny

We have thus seen how phage replication is tolerated, i.e., how a reliable response to the initial MOI is achieved despite the subsequent change in viral copy number. Recall, however, that comparing CI-Cro trajectories in replicating (**Figure 3B**) and nonreplicating (**Figure 3A**) phages indicated that viral replication is not only tolerated but, in fact, required for the existence of an MOI-dependent lysis-to-lysogeny transition. To understand why that is the case, we first addressed the absence of a lytic choice following *P*- infection at MOI = 1. Our model indicates that, in both *P*+ and *P*-, *cro* transcription from P_R_ (per phage) is repressed >2-fold within ~10 minutes of infection (**Figure 6A**; **Figure S14** depicts the corresponding experimental data). However, the presence of additional gene copies in the *P*+ case results in higher Cro concentration later in the infection, sufficient to cross the lytic threshold (**Figure 6B**). Viral replication thus serves to boost total Cro expression despite repression of its transcription at the individual phage level.

As for the effect of replication on the lysogenic choice, we find that *cI* transcription from P_RE_ during *P*- infection follows a similar pattern to *P*+, namely, a single pulse whose duration and amplitude per phage depend only weakly on MOI (**Figure S15**, compare to **Figures 4A** and **S10**). However, as in the case of Cro, the presence of added gene copies during *P*+ infection leads to considerably higher CI accumulation than for *P*- (**Figure 6C**). Consequently, while replicating phages reach the lysogenic CI threshold at MOI = 2, nonreplicating ones require a higher MOI to reach that threshold and establish lysogeny. The theoretical prediction that *P*- phage lysogenizes at higher MOI than *P*+ is born out in experiments (Kourilsky (1973) and **Figure 6D**).

To generalize the effect of viral replication on the lysis/lysogeny decision, we simulated infections at MOI in the range 1–7, while varying the phage replication rate from zero to 1.5x that of *P*+ phage. Determining the infection outcome at each MOI and replication rate yielded the twodimensional “fate diagram” shown in **Figure 6E**. Consistent with the discussion above, we find that both the existence of a lytic outcome and the minimal MOI at which the transition to lysogeny occurs depend on the viral replication rate. The ability to replicate is not by itself sufficient to enable the lytic pathway. Rather, there is a minimum required replication rate, below which, MOI = 1 infections fail to achieve either outcome. As for lysogeny, the MOI at which this fate is chosen decreases with the viral replication rate (**Figure 6E**). When replication is sufficiently rapid, the model predicts a lysogenic outcome even at MOI = 1. While we are unaware of an experimental test for this prediction, we note that the model predicts a similar behavior when the CII activity window is extended by inhibition of CII degradation, a result validated by experiments (Kobiler et al., 2007)(**Figure 6D** and **Figure S16**).

Interestingly, there are two regions where the system’s trajectories cross both the lytic and lysogenic thresholds in the course of infection (**Figure 6E**). The first of those occurs at high replication rates, in the MOI range between lysis and lysogeny. We are uncertain how to interpret this feature, but it is intriguing to note that, at the single-cell level, it may correspond to the range of infection parameters where stochastic effects become important and, consequently, individual cells exhibit different fates, as reported experimentally (Lieb, 1953; St-Pierre and Endy, 2008; Zeng et al., 2010). The analysis of cellular heterogeneity is outside the premise of our current model, which captures the population-averaged behavior only. A second region where mixed outcomes are predicted is found for MOI ≳ 5, and may correspond to a scenario where the overexpression of viral proteins results in the halting of cell growth, rather than a lytic or lysogenic outcome, consistent with experimental data (Zeng et al., 2010).

## DISCUSSION

The combination of single-molecule genome and mRNA measurement in individual cells with theoretical modeling provided us with new insights into the way lambda counts coinfecting phages to bias the lysis-lysogeny decision. Early theoretical studies of the post-infection decision sought insight to the phage’s binary choice in the toggle switch comprised of the mutually antagonistic CI and Cro, which governs lysogenic maintenance and lytic induction (Avlund et al., 2009; Reinitz and Vaisnys, 1990; Shea and Ackers, 1985; Weitz et al., 2008). This famous “genetic switch” exhibits bistability (Bednarz et al., 2014), with well-defined states characterized by high CI (lysogeny) and high Cro (lytic onset), respectively. These features, and the tremendous body of experimental and theoretical knowledge that has accrued about the pairwise CI/Cro interactions (Court et al., 2007; Ptashne, 2004), explain the focus on this element as the key to the decision. Moreover, theoretical work has shown that MOI could indeed drive a bifurcation of the CI/Cro switch’s steady state, consistent with a transition from lysis to lysogeny (Avlund et al., 2009; Weitz et al., 2008). Our analysis above, however, suggests that this is not how the lambda decision unfolds. Instead, each phage initially attempts to execute a preset pattern of gene expression, independent of the MOI. In this cascade of events, *cI* transcription takes place predominantly through the transient activation of P_RE_ by CII, while *cI* autoregulatory expression from P_RM_ is unnecessary. The resulting cellular CI expression is approximately linear in MOI, indicating the absence of an ultrasensitive response to CII as previously suggested (Cortes et al., 2017; Kobiler et al., 2005; Vohradsky, 2001). Given this fixed gene expression cascade at the individual phage level, it is the introduction of time-varying gene dosage due to viral replication—rather than the standalone topology of the viral circuit—that enables the subsequent divergence of gene expression trajectories and cell-fate choices at low and high MOI.

The time-varying viral copy number is found to be critical for both possible outcomes of infection. The choice of lysogeny depends on the number of lambda copies present during the early decision window, set by CII activation of P_RE_ (**Figure 4A** above). The finite response window immunizes the lambda decision to changes in viral copy number that take place outside it, thus allowing reliable detection of the initial MOI. However, the role of viral copy number does not end then, and is, in fact, crucial for reaching the protein threshold required for establishing either fate: Cro for lysis, or CI for lysogeny. Cro (and by proxy, the lytic activator Q) continues to accumulate through late infection, reaching the lytic decision threshold ~50 minutes post-infection (**Figure 6B**). Cro’s continued accumulation is driven by late viral replication, despite the repression of P_R_ at the single phage level (**Figure 6A**). As for lysogeny, while this fate is achievable in the absence of viral replication, it is replication that ensures that co-infection by more than one phage is sufficient to drive lysogeny; in the absence of replication, CI accumulates insufficiently, and higher MOI is required to reach the lysogenic threshold (**Figure 6C**).

The relation between gene dosage and the output of genetic networks has been explored in diverse biological contexts, both natural (Guo et al., 1996; Narula et al., 2015; Pollack et al., 2002) and synthetic (Baumgart et al., 2017; Becskei and Serrano, 2000; Bleris et al., 2011). In some instances, changes in gene copy number were found to have a significant effect on phenotype (Cook et al., 2012; Maron et al., 2013; Wright et al., 2009) whereas in other cases, mechanisms of dosage compensation buffer the network output from such changes (Acar et al., 2010; Hose et al., 2015; Segall-Shapiro et al., 2018; Voichek et al., 2016). The lambda decision exhibits a richness of dosage response beyond what was previously documented, with the transcriptional output either linear in dosage, or partially compensating it, for different genes in the network, at different times during the infection. Consequently, viral replication is found to facilitate, rather than hinder, the implementation of a reliable decision. Since cellular decisions frequently take place even as gene copy number is changing (Desponds et al., 2020; Hwang et al., 2017; Weinberger, 2015), this inextricable coupling of gene dosage and network output cannot be ignored if one aims for a predictive description of cellular decision making.

## Acknowledgements

We are grateful to the following people for their generous advice and for providing reagents: R. Arbel-Goren, S. Austin, G. Balászi, M. Cortes, I. Dodd, M. Feiss, J. Harris, C. Hayter, K. Shearwin, J. Stavans, F. St-Pierre, L. Thomason, G. Vasen, J. Weitz, L. Weinberger, Z. Yu, L. Zeng, and all members of the Golding and Igoshin groups. This work was supported by the National Science Foundation Center for Theoretical Biological Physics (NSF PHY-2019745) and grant PHY-1522550. The work was supported in part by the Big-Data Private-Cloud Research Cyberinfrastructure MRI-award funded by NSF under grant CNS-1338099 and by Rice University. Work in the Golding lab is supported by the National Institutes of Health grant R01 GM082837 and the National Science Foundation grant PHY 1430124. Igoshin also acknowledges support by the Welch Foundation Grant C-1995 and the National Science Foundation grant MCB-1616755.

## Supplementary Information

Experimental Methods

Theoretical Methods

Supplementary Figures 1–16

Supplementary Tables 1–6

## Supplementary Information

### Experimental Methods

#### 1 Growth media and conditions

##### 1.1 Media

Unless otherwise noted, the growth medium for all *E. coli* strains was LB (Lennox recipe [1]): One liter of medium was prepared with 10 g tryptone (BD Biosciences), 5 g yeast extract (BD Biosciences), and 5 g NaCl (Fisher Scientific), pH adjusted using 1 μM NaOH (Fisher Scientific). When applicable, the LB medium was supplemented with 10 mM MgSO_4_ (Fisher Scientific) and 0.2% maltose (Fisher Scientific), hereafter denoted as LBMM, or with 10 mM MgSO_4_ and 0.2% glucose (Fisher Scientific), hereafter denoted as LBGM. Phage plaque assays (see below) were performed using NZYM agar. One liter of NZYM medium was prepared with 22 g NZYM media (Teknova) and adjusted using 10 μM NaOH. LB and NZYM agar plates were prepared using the media above, with 1.5% weight/volume agar (BD Biosciences). All media—with and without agar—were autoclaved using a liquid cycle (121°C, at least 25 minutes) for sterilization.

##### 1.2 Growth conditions

Cells were streaked and grown overnight (14–16 hours) on LB agar plates supplemented with antibiotics when applicable: 100 μg/mL ampicillin (Fisher Scientific), 50 μg/mL kanamycin (Fisher Scientific). Plates were incubated at the appropriate temperature: 30°C for temperature-sensitive lysogens, 37°C otherwise. From plates, fresh colonies were inoculated in 2 mL of LB or LBMM in 14 mL round-bottom test tubes (Falcon), supplemented with antibiotics when applicable. The overnight cultures were grown for 14–16 hours at the appropriate temperature with aeration (220 rpm) in a MaxQ 4000 Benchtop Orbital Shaker (Thermo Scientific). The growth conditions of overday cultures are described later for each specific experiment (**Sections 3, 4 and 9**). When applicable, the overday cultures were also supplemented with isopropyl-β- thiogalactoside (IPTG, Sigma-Aldrich) at the required concentration to induce ParB production. The overday cultures were grown in a MaxQ 7000 Water Bath Orbital Shaker (Thermo Scientific).

#### 2 Bacterial strains and plasmids

All bacterial and phage strains, plasmids, primers and DNA oligos are listed in **Tables S1–S5**. We used *E. coli* strains LE392 and MG1655. The phage used, λ_TY11_ (*λ cI*857 *Pam*80 *stf::P1parS-kanR*, described in **Section 3** below), carries an amber mutation in the *P* gene [2], which allows replication of the phage genome in the amber suppressor strain LE392 but not in the wild type strain MG1655 [3]. We therefore utilized LE392 during construction and propagation of the phage (**Section 3**), and MG1655 for infection experiments where viral copy numbers were to be held constant (**Sections 4 and 9**). Intracellular phage genomes were labelled using the ParB-*parS* system [4, 5]. The *parS* sequence of phage P1, engineered into the lambda genome, was bound by a fluorescent fusion version of the P1 ParB protein (CFP-ParB) expressed from the plasmid pALA3047 (gift of Stuart Austin).

#### 3 Phage construction

To fluorescently label the phage genome in the cell, we inserted the *parS* sequence and a kanamycin resistance cassette to the *stf* region of λ *Pam*80 (gift of Lynn Thomason). Next, we converted the wild-type *cI* in the phage’s genome to a temperature-sensitive allele (*cI*857), to allow prophage induction via temperature shift [3] and for consistency with previous studies [5–8]. The details of phage construction are provided in the following subsections.

##### 3.1 Construction of the pTY001 plasmid containing the *parS* sequence

We first constructed the plasmid pTY001, which is used to insert the *parS* sequence into the phage genomes, as follows. The plasmid carries a cassette with *parS* and *kanR* sequences, flanked by homologies to lambda *stf* region (considered nonessential for phages derived from lambda PaPa [9]). Our template was a phage strain in which the *kanR-parS* cassette was placed within the *stf* region (gift of Joel Stavans) [5]. We first amplified the *kanR-parS* cassette (with *stf* homologies) using the primers *p1-parS-FP* and *p1-parS-RP* (**Table S4**). Then, we cut both the PCR product and a pBS-SK vector (Stratagene) using KpnI and SacII (New England Biolabs). The insert and the vector backbone were then purified by electrophoresis using 1% agarose gel (Bio-Rad), followed by DNA extraction from the gel using Wizard SV Gel and PCR Clean-Up System (Promega). Finally, the insert and the vector were ligated using T4 DNA ligase (New England Biolabs). The pTY001 plasmid was then transformed into strain LE392 using the Bio-Rad MicroPulser Electroporator in accordance with the instrument’s instruction manual.

##### 3.2 Construction of the λ_TY8_ phage containing the *parS* sequence

We next constructed phage λ_TY8_ (*λ Pam*80 *stf::P1parS-kanR*) by performing a phage-by-plasmid cross [10] between the parental phage λ *Pam*80 (carrying a C-to-T mutation at the 39,759 position in the lambda genome [2], an amber mutation in the *P* gene) and the plasmid pTY001 (described above), as follows. An overday culture of the host LE392 carrying plasmid pTY001 was prepared by diluting the overnight culture 1:1000 into 110 mL LBMM. The overday culture was grown in a 1 L baffled Erlenmeyer flask at 37°C with 220 rpm aeration. Upon reaching OD_600_ ≈ 0.4 (measured using the Bio-Rad SmartSpec Plus Spectrophotometer), 20 mL of culture (containing approximately 2×10^9^ cells) were centrifuged at 4,000×g for 10 minutes at 4°C (using the Thermo Scientific Sorvall Legend XTR Centrifuge), and the supernatant was removed. The cells were then resuspended in 200 μL of fresh LBMM medium. For infection, 200 μL of phage λ *Pam*80 was mixed 1:1 (equal volume) with the concentrated cells in a 1.5 mL Eppendorf tube, to reach MOI ≈ 0.5. The infection mixture was incubated at 37°C for 15 minutes, then diluted into 15 mL of LBGM (prewarmed to 37°C) in a 125 mL baffled Erlenmeyer flask and shaken at 37°C for 120 minutes to complete a lytic cycle. Chloroform (final concentration of 5%) was then added to the media to lyse all remaining cells. The solution was centrifuged at 2,000×g for 10 minutes at 4°C to remove the cell debris, and the phage stock (containing both the unmodified λ *Pam*80 and the modified λ *Pam*80 *stf::P1parS-kanR*, named λ_TY8_) was harvested.

To screen for the recombinant phage λ_TY8_, we first performed lysogenization and selected for lysogens harboring kanamycin resistance, as follows. An overday culture of LE392 was prepared by diluting the overnight culture 1:1000 into 25 mL of LBMM, and the culture was grown in a 250 mL baffled Erlenmeyer flask at 37°C with 220 rpm aeration. Upon reaching OD_600_ ≈ 0.4, cells were harvested by centrifugation at 4,000×g for 10 minutes at 4°C. The supernatant was removed, and cells were resuspended in 2.5 mL of fresh LBMM (at approximately 1×10^9^ cells/mL). The phage stock above, containing λ_TY8_ as well as non-recombinant phages, was diluted in SM buffer (Teknova) to reach a concentration of approximately 1×10^9^ plaque-forming units (PFU)/mL, then mixed with the concentrated cells to reach MOI ≈ 1. The infection mixture was incubated at room temperature for 20 minutes, then diluted into 1 mL of LBGM (prewarmed to 30°C) in 14 mL round-bottom test tubes (Falcon) and shaken at 30°C for 1.5 hours. 100 μL of the culture was plated on LB agar plates supplemented with 50 μg/mL kanamycin. The plates were incubated overnight at 30°C. Only lysogens harboring recombinant prophages, which carry the kanamycin-resistance cassette, survived and formed colonies under this selection.

Poly-lysogenic cells, carrying multiple tandem prophages, may harbor a mixture of nonrecombinant and recombined prophages [11], with the latter conferring kanamycin resistance to the cell. To screen for monolysogens, we picked a number of colonies from the lysogens above and performed PCR using primers *e.coli_attB, lambda_attB* and *lambda_int* (**Table S4**) to detect for the presence (or absence) of the junction between multiple prophages within a polylysogen [12]. Next, lysogens carrying a single recombinant prophage (denoted λ_TY8_) underwent prophage induction using mitomycin C (Fisher Scientific) following the protocol of [3], and the phage lysate containing λ_TY8_ was harvested.

##### 3.3 Construction of the temperature-sensitive phage λ_TY11_

We next constructed λ_TY11_ (*λ cI*857 *Pam80 stf::P1-parS-kanR*) by converting the *cI^wt^* allele in λ_TY8_ (described above) to the temperature-sensitive allele *cI*857 (consisting of a C-to-T mutation at the 37,742 position in the lambda genome [3]). The phage genotype was modified using a recombineering protocol for modifying an infecting phage (here, λ_TY8_) [13]. As host we used strain LE392 carrying the plasmid pKM208 (Addgene). This plasmid contains the lambda *gam*, *beta* and *exo* genes [14], required for lambda recombination [15]. We designed the single-strand oligos *cIts-oligo-R* and *cIts-oligo-F* (**Table S4**), which contained the target single-nucleotide mutation flanked by around 40 nt homologous sequences at both sides. We then followed the protocol of [13] and harvested the phage lysate, which contained both the unmodified λ_TY8_ and the recombinant λ_TY11_. To screen for the recombinant phage λ_TY11_, we took advantage of the fact that, at 37°C, the phage would form clear plaques due to its temperature-sensitive *cI*857 allele, whereas the unmodified λ_TY8_ would form turbid plaques [3, 16]. We first diluted the phage lysate into SM buffer (Teknova), then mixed 10 μL of the diluted phage lysate (contains approximately 5×10^3^ phages) with 100 μL of LE392 cells (at approximately 1×10^9^ cells/mL) in a 1.5 mL Eppendorf tube. The mixture was incubated at 37°C for 15 minutes, then added to 3 mL of molten 0.7% NZYM agar (maintained at 47°C) in a 14 mL round-bottom test tube (Falcon). The mixture was then poured onto a 1.5% NZYM agar plate and left to solidify at room temperature for approximately 15 minutes. The plate was incubated overnight at 37°C to allow for plaque formation.

The next day, ~0.1% of plaques were clear, indicating successful conversion to the temperature-sensitive allele. To collect phages from the clear plaques, we first picked a whole plaque by penetrating the top agar around the clear plaque using a 100 μL pipet tip with a wide-cut end. Then, the agar piece with the plaque was soaked in 100 μL SM buffer in a 1.5 mL Eppendorf tube, yielding a lysate of the recombinant phages. Using a lysogenization protocol similar to **Section 3.2** above, we generated the lysogenic strain LE392 λ_TY11_, for storage. The genotypes of *cI*857 and *Pam*80 in the recombinant prophage were confirmed by sequencing (Lone Star Labs Genetic Sequencing) using the primers *cI-seq-F, cI-seq-R, P-seq-F* and *P-seq-R* (**Table S4**). The LE392 λ_TY11_ lysogen underwent prophage induction at 42°C following the protocol of [3], to obtain a phage lysate of λ_TY11_.

#### 4 Phage infection

##### 4.1 Bulk lysogenization assay

We measured the probability of lysogenization as a function of MOI, using a protocol adapted from [7, 17]. An overnight culture of the host was diluted 1:1000 into 25 mL LBMM and grown in a 125 mL baffled Erlenmeyer flask at 37°C with 220 rpm aeration. The phage lysate was diluted in SM buffer (Teknova) to yield a 4-fold dilution series between 10^7^ and 10^11^ PFU/mL. Upon reaching OD_600_ ≈ 0.4, cells were centrifuged at 1,000×g for 10 minutes at 4°C, and the supernatant was removed by pipetting. The cell pellet was resuspended in 1 mL of fresh, ice-cold LBMM. Then, 20 μL of the concentrated bacteria was combined with an equal volume of the diluted phage solution (at different concentrations, measured by standard plaque assay [18]), resulting in MOI in a range of ≈ 0.01-100. The infection mixture was incubated on ice for 30 minutes, followed by an additional 5-minute incubation in a 35°C water bath to trigger injection of the phage DNA [7]. Next, we diluted 10 μL of each infection mixture into 1 mL LBGM (prewarmed at 30°C) in 14 mL round-bottom test tubes (Falcon), and incubated the diluted mixtures at 30°C for 45 minutes with 220 rpm aeration. The cells were then diluted in ice-cold 1×PBS buffer to create a 10-fold dilution series between 10^3^ and 10^7^ cells/mL, and 100 μL of the diluted cells were plated on LB agar plates supplemented with 50 μg/mL kanamycin. The uninfected cells were also diluted and plated in a similar manner on LB agar plates. All plates were incubated overnight at 32°C. The lysogenization frequency was determined by dividing the number of lysogen colonies (on kanamycin-selective plates) and uninfected colonies (on non-selective plates) on the next day.

##### 4.2 Infection followed by smFISH

An overnight culture of MG1655 carrying plasmid pALA3047 was diluted 1:1000 into 75 mL of LBMM supplemented with 10 μM IPTG and grown in a 500 mL baffled Erlenmeyer flask at 37°C with 220 rpm aeration. Upon reaching OD_600_ ≈ 0.4, cells were transferred to 50 mL centrifuge tubes (Corning) and centrifuged at 1000×g for 10 minutes at 4°C. The supernatant was carefully decanted, and the cells were resuspended in fresh, ice-cold LBMM supplemented with 10 μM IPTG at 100× the original concentration (approximately 7×10^9^ cells/mL after resuspension). We left some host cells uninfected as the negative control, treating them according to the smFISH procedure described in **Section 5** below. Next, 500 μL of the concentrated host cells were mixed with 70 μL of λ_TY11_ phage lysate (prepared as described in **Section 3.3**) to reach MOI ≈ 2. The infection mixture was incubated on ice for 30 minutes, followed by an additional 5 minute incubation in 35°C water bath to trigger injection of the phage DNA [7]. Next, the infection mixture was diluted 1:1000 into 400 mL of LBGM (prewarmed to 30°C) supplemented with 10 μM IPTG, split into two 2 L baffled Erlenmeyer flasks, and shaken at 30°C. At various time points, 30 mL of the culture was collected and treated according to the smFISH procedure described in **Section 5** below.

##### 4.3 Infection followed by DNA extraction and qPCR

The crude lysates of phage λ *cI*857 *Sam*7 (for the λ *P*+ data series) and of phage λ *Pam*80 (for the λ *P*- data series) were produced by heat induction as in [3] from lysogenic cells (gift of Mike Feiss and Lynn Thomason, respectively). For λ *P*+, the *Sam*7 genotype was selected because the amber mutation in the phage *S* gene prevents cell lysis in the non-suppressor MG1655 strain [3]. This allowed us to measure the number of phages in the cells throughout the lytic cycle. An overnight culture of MG1655 was diluted 1:1000 into 100 mL of LBMM in a 1 L baffled Erlenmeyer flask and grown at 37°C with 220 rpm aeration. Upon reaching OD_600_ ≈ 0.4, cells were centrifuged at 2,000×g for 10 minutes at 4°C. The supernatant was removed, and the cells were resuspended in fresh, ice-cold LBMM at 100× the original concentration (approximately 7×10^9^ cells/mL). The phage lysate was diluted in ice-cold SM buffer (Teknova) to achieve a titer of approximately 4×10^9^ PFU/mL. Next, 300 μL of the cold phage lysate was added to 300 μL of the cold concentrated bacteria in a 1.5 mL Eppendorf tube to reach MOI ≈ 0.5. The infection mixture was gently mixed by pipetting. A negative control was also prepared by mixing cells with DEPC-H_2_O (Invitrogen), using the same 1:1 volume ratio. The infection mixture and the negative control were incubated on ice for 30 minutes to allow phage adsorption. Genome injection was triggered by shifting the samples to a 35°C water bath for 5 minutes [7]. Then, 50 μL aliquots of the infection mixture (or the negative control) were each diluted 1:500 into 25 mL of LBGM (prewarmed to 30°C) in multiple 250 mL baffled Erlenmeyer flasks, then grown at 30°C and 220 rpm aeration. At each time point, the entire diluted infection mixture from single flasks was treated according to the DNA extraction procedure described in **Section 9** below.

#### 5 Single-molecule fluorescence *in situ* hybridization (smFISH)

The smFISH protocol was described in detail previously [19]. Briefly, sets of antisense DNA oligo probes were designed against lambda *cI*, *cro* and *cII* mRNA, synthesized with a 3’ amine modification (LGC Biosearch Technologies), and all oligos against a given gene were pooled together. We covalently linked the *cI*, *cro* and *cII* probe sets to TAMRA, Alexa 594 and Alexa 647 (Invitrogen) respectively, and purified them using ethanol precipitation. The probe concentration and dye labeling efficiency were measured using a NanoDrop 2000 spectrophotometer (Thermo Scientific). Probe sequences are listed in **Table S5**. Following the cell growth and infection procedure described in **Section 4.2** above, cells were fixed and permeabilized, then incubated with the fluorescently labeled probe sets, washed, and finally imaged as described in **Section 6** below. We made the following modifications relative to the original protocol from [19]. At each time point, 30 mL cell culture was directly mixed with 7.5 mL of 18.5% formaldehyde solution in 5×PBS (to a final concentration of 3.7% formaldehyde in 1×PBS) and nutated for 30 minutes at room temperature. In addition, when permeabilizing the cells, we resuspended the cell pellets in 750 μL of DEPC-H_2_O (Invitrogen), mixed thoroughly with 250 μL of 100% ethanol by pipetting, then mixed gently for 1 hour at room temperature using a nutator. We found that a final ethanol concentration of 25%, instead of 70% as in the original protocol, better preserved the fluorescence signal of CFP-ParB without harming the permeability of cells to smFISH probes (Data not shown).

#### 6 Microscopy

We used an inverted epifluorescence microscope (Eclipse T*i*, Nikon), equipped with motorized stage control (ProScan III, Prior Scientific), a universal specimen holder, a mercury lamp (Intensilight C-HGFIE, Nikon), and a CMOS camera (Prime 95B, Photometrics). A ×100, NA 1.40, oil-immersion phase-contrast objective (Plan Apo, Nikon) was used, as well as a ×2.5 magnification lens (Nikon) in front of the camera. The fluorescent filter sets used in the study were as follows: CFP (Nikon, 96341), Narrow Cy3 (Chroma, SP102v1), Narrow Cy5 (Chroma, 49307), and a customized set for imaging Alexa 594 (Omega, excitation filter: 590± 10 nm; dichroic beam splitter: 610 nm; emission filter: 630±30 nm).

After the fixation, hybridization and washing steps (described in **Section 5**), we mounted the cells between two coverslips as described in [19]. The sample was then placed onto the microscope’s slide holder and the cells were visually located using the phase-contrast channel. Images were acquired in the following order: phase-contrast (100 ms, to detect the cell outline), CFP (200 ms, CFP-ParB), Narrow Cy3 (500 ms, *cI*-TAMRA), 594 cube (500 ms, *cro*-Alexa 594), and Narrow Cy5 (500 ms, *cII*-Alexa 647). Snapshots were taken at 5 z-positions (focal planes) with steps of 300 nm. A set of images with multiple z-positions is denoted as an “image stack” and the image of each z position as a “z-slice”. Images were acquired at multiple positions on the slide, to image a total of 400-2000 cells per sample (typically 9-20 positions).

#### 7 Cell segmentation and spot recognition

##### 7.1 Cell segmentation

Our procedure for identifying cells from the phase-contrast channel follows [19, 20]. In every image stack, we first identified the “in-focus” z-slice, defined to be the one with the largest variance among pixels. Then we used Schnitzcells [21] to generate cell segmentation masks, i.e., matrices with the same dimension as the phase-contrast image, in which pixels of cells have integer values corresponding to the identification number of the cell, while non-cell pixels have a value of zero. Finally, we visually inspected the segmentation results; poorly segmented cells were either manually corrected or discarded using the graphical user interface within the software.

##### 7.2 Spot recognition

Following [19, 20], we used Spätzcells [19] to identify and measure the properties of fluorescent foci (spots) from the different fluorescent channels. Briefly, the Spätzcells software first identified spots by finding two-dimensional local maxima in the fluorescence intensity above a user-defined “detection threshold”. Next, the fluorescence intensity profile around each spot was fitted to a 2D elliptical Gaussian. The properties of each spot were obtained from the fitting results, including the position, spot area, peak height of the fitted Gaussian, and spot intensity (integration of the volume underneath the fitted Gaussian). In cases where other spots were close to the spot being fitted, the software performed a 2D multi-Gaussian fit over all detected spots instead.

##### 7.3 Discarding false-positive spots

Using a low detection threshold during spot recognition (**Section 7.2**) ensured that all genuine spots were detected. However, because of the low threshold, the number of false positives (from background noise, or nonspecific binding by smFISH probes and CFP-Par) also increased. To discard the false positives, we performed a gating procedure following [19, 20]. Briefly, we compared the 2D scatter plots of peak height versus spot area for all detected spots in the experimental samples (infected cells) to that from the negative control (uninfected host cells). A polygon was manually chosen in the plane of peak height and spot area, such that most spots from the negative sample were located outside of it. This polygon served as the gating criteria for all samples, with spots located outside of discarded. The choice of gating was confirmed by manual inspection of spots in a subset of images.

#### 8 Data analysis following cell segmentation and spot recognition

##### 8.1 mRNA quantification

Defining the fluorescence intensity of a single mRNA molecule was performed as described in [19, 20]. Briefly, after discarding the false positive spots, we examined the early infection samples, which exhibit low mRNA levels, and in which individual mRNA molecules were spatially separated. The histograms of spot intensities were fitted to a sum of three Gaussians corresponding to one, two, and three mRNA molecules per spot. The fluorescence intensity of a single mRNA molecule was estimated to be the center of the first Gaussian. We then divided the measured spot intensity of each mRNA spot by the single-mRNA intensity to obtain the number of mRNA molecule number in that spot. The total number of mRNA molecules in a given cell was calculated by summing the mRNA molecule numbers represented by all spots within the cell.

##### 8.2 Identification of single-cell MOI

To verify that CFP-ParB spots correspond to individual phage genomes, we confirmed that, in lysogenic cells, the mean number of CFP-ParB spots per cell was consistent with the expected prophage copy-number under the specific growth conditions [22, 23] (**Figure S2**). Therefore, our estimated number of infecting phages in each individual cell (single-cell MOI) was the CFP-ParB spot number in the cell, as measured by Spätzcells (**Section 7.2**). Infected cells were grouped based on single-cell MOI and the corresponding levels of *cI*, *cro* and *cII* mRNA calculated for each group were used in constraining the mathematical model, as described in **Theoretical Methods**.

#### 9 DNA extraction and quantitative PCR (qPCR)

##### 9.1 Calibration curves

We used two pairs of primers: One targeting a 154-bp region in the *cI* gene of phage lambda, and another targeting a 150-bp region in *lacZ* of *E. coli* [24]. The primer sequences are provided in **Table S4**. The amplification efficiencies of the primer sets were first determined as follows. Two 25 mL cultures of non-lysogens (MG1655, containing *lacZ*) and lysogens (MG1655 λ_IG2903_, containing both *cI* and *lacZ*) were grown at 30°C in LB supplemented with 10 mM MgSO_4_ with 220 rpm aeration. Upon reaching OD_600_ ≈ 0.4, 2 mL of cells (containing approximately 2×10^8^ cells) were centrifuged at 21,130×g (max speed) for 1 minute using a Eppendorf 5420 centrifuge, and the supernatant was removed. Genomic DNA was extracted from the cell pellet using the PureLink Genomic DNA Mini Kit (Invitrogen), according to the kit’s instructions for Gramnegative bacteria. The concentration and quality (A260/A280 and A260/A230) of the DNA extracts were measured using a NanoDrop 2000 spectrophotometer (Thermo Scientific). A serial dilution series of the lysogen DNA was prepared, spanning five orders of magnitude (5 pg to 50 ng of DNA per 20 μL reaction). The DNA extracted from non-lysogens served as the negative control for the *cI* target. Each DNA sample was then subjected to both pairs of primers (for *cI* and *lacZ* targets). qPCR was performed using SsoAdvanced Universal SYBR Green Supermix (Bio-Rad), in either a MiniOpticon Real-Time PCR System (Bio-Rad) or a CFX Connect Real-Time PCR Detection System (Bio-Rad). We prepared 20 μL of each qPCR reaction comprising 1× supermix, 500 nM of the forward and reverse primers (each), DNA template (5 pg to 50 ng, diluted as above), and DEPC-H_2_O. No-template controls (NTC) were also prepared by replacing DNA with DEPC-H_2_O. Each sample was prepared in technical duplicates or triplicates. The thermal cycle was chosen based the instruction manual, specifically, 3 minutes at 98°C, followed by 40 cycles of 15 seconds at 98°C and 30 seconds at 60°C, then finally a melt curve analysis from 65°C to 95°C. The amplification efficiencies of the *cI* and *lacZ* targets were determined from the calibration curves following standard procedure [25, 26]. Calculated from 10 and 8 biological replicates, respectively, the efficiencies for the *cI* and *lacZ* targets were 1.998 (or 99.8%) and 1.956 (or 95.6%), respectively.

##### 9.2 Quantification of phage copy numbers following infection

Infection of MG1655 by λ *cI*857 *Sam*7 or λ *Pam*80 was performed as described in **Section 4** above. At each time point, 25 mL of the diluted infection mixture (containing approximately 2×10^8^ cells) was poured into a pre-chilled 50 mL centrifuge tube (Corning) and incubated in an iced water bath to stop cell activity. The negative control (cells mixed with DEPC-H_2_O, no phage) was collected at the end of the experiment. All samples were centrifuged at 4,000×g for 10 minutes at 4°C, and DNA was extracted from the cell pellets, followed by concentration and quality measurements as above. The qPCR reactions were set up and performed as above, with 50 pg of template DNA for each sample. Using the target-specific amplification efficiency (measured above), we performed a relative quantification of *cI*, with *lacZ* serving as a reference gene in accordance with the efficiency-correction quantification method [25, 26], yielding the ratio of *cI* to *lacZ* templates in each sample. We then used these values and the estimated copy number of the *lacZ* locus per cell (approximately 2.6 under the relevant growth conditions [22]) to estimate the average copy number of the *cI* locus—a proxy for the number of lambda genomes per cell in each sample.

## Theoretical Methods

### 1 Overview of the Governing Differential Equations

As discussed in the main text, our model focuses on three genes at the center of the lambda cellfate decision: *cI, cro*, and *cII*. Based on the known interactions in the lambda network (discussed in the main text and in more detail below in **Section 2**), we constructed a deterministic model describing the dynamics of *cI, cro*, and *cII* mRNA and protein concentrations, as well as viral concentration. The interactions within this 3-gene network are shown schematically in **Figure 2A** (main text). Denoting the mRNA and protein concentrations of gene *x* as [*m_x_*] and [X] respectively, and viral concentration as [λ], the model consists of the following differential equations:

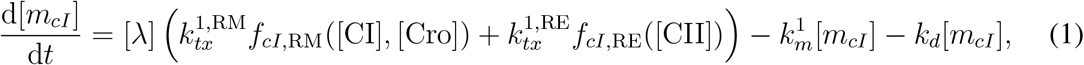

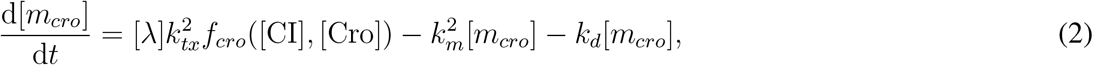

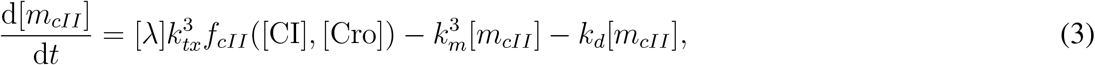

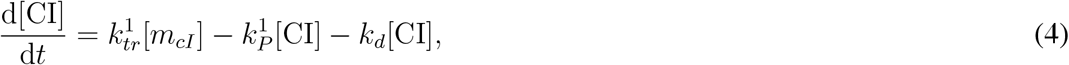

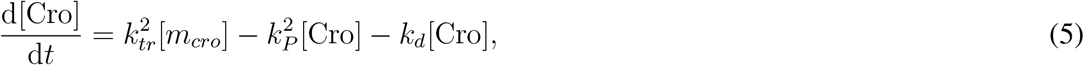

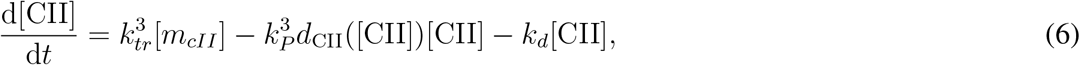

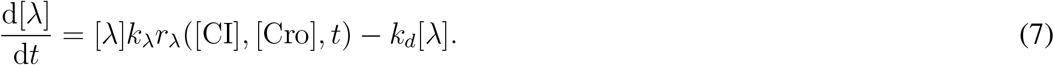

Here, 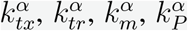 denote the rate constants for transcription, translation, mRNA degradation, and protein degradation respectively (with superscript *α* denoting the affiliated gene: 1 = *cI*, 2 = *cro*, and 3 = *cII*), *k*_λ_ is the rate of viral replication, and *k_d_* is the rate of dilution due to cell growth. As cI is transcribed from two promoters (as discussed in the main text and **Section 2.1.1** below), we use additional RM and RE superscripts to differentiate transcription rates from each promoter. Values of these and other parameters used to generate the figures in the main text are estimated as described in **Section 4** and summarized in **Theoretical Methods Table 3 (Section 7**).

Equations 1–7 are based on several simplifying assumptions commonly used for modeling lambda and other bacterial systems. Following previous works in lambda [1, 2], we assumed that transcription rates are proportional to viral concentration, i.e. that transcription fluxes from each additional viral copy are the same. Following standard practice [3], we modeled translation as being proportional to mRNA concentration in Eqs. 4–6, and modeled mRNA degradation (Eqs. 1–3) as a first-order reaction. Assuming exponential growth of cell volume (following previous models of lambda [4, 5] and of other bacterial systems [6–8]) leads to the first-order effective degradation term in Eqs. 1–7 with rate *k_d_*. To model transcriptional regulation in Eqs. 1–3, we introduced transcription regulatory functions (*f_cro_, *f*_cll_, f_ci,RM_*, and *f_cI_*,_RE_). These functions are dependent on transcription factor concentrations and used as multiplicative factors to change the transcription rate of individual genes [9]. To describe protein degradation, we modeled nonspecific decay of the stable proteins CI (Eq. 4) and Cro (Eq. 5) as a first-order reaction with a constant rate (again in line with common practice [3]). In contrast, CII is actively degraded, and its degradation is regulated [10]. Therefore, the degradation term in Eq. 6 is multiplied by a degradation regulatory function, *d*_CII_. Viral replication in Eq. 7 is modeled as exponential growth of the number of viruses with the rate proportional to a replication regulatory function (r_λ_). The functional forms and biological basis for all regulatory functions are described in **Section 2**.

All codes and parameter/datasets can be found in the following GitHub repository: https://github.com/sethtcoleman/Replication-Manuscript.git

#### 2 Formulation of Regulatory Functions

Although phage lambda is a comparatively well-characterized system, the molecular details of many regulatory interactions are still uncertain [11]. We therefore designed our model to capture known regulatory interactions phenomenologically. Below we describe how transcription (*f_cro_*, *f_cll_*, *f_ci_*,_RM_, and *f_cI_*,_RE_), CII degradation (*d*_CII_), and viral replication (*r*_λ_) regulatory functions were formulated.

##### 2.1 Transcription

Motivated by previous work that characterized transcription regulation phenomenologically [12, 13], we modeled the effect of transcription factors on promoter activity at the single copy level using functions composed of standard Hill terms for activation ([X]*^n^*/ (*K^n^* + [X]*^n^*)), repression (*K^n^*/ (*K^n^* + [X]*^n^*)), and their generalizations to combinatorial gene regulation by multiple factors [9]. We applied this approach to *cI* transcription from P_RM_ and P_RE_ and *cro* and *cII* transcription from P_R_.

###### 2.1.1 *cI* Transcription

cI is transcribed from two promoters: P_RE_ (CII-activated) and P_RM_ (regulated by CI and Cro). For modeling *cI* transcription from P_RE_, we followed common practice and ignored basal transcription [1,2,4]. The single phage transcription regulatory function for cI expression from P_RE_ is a Hill function

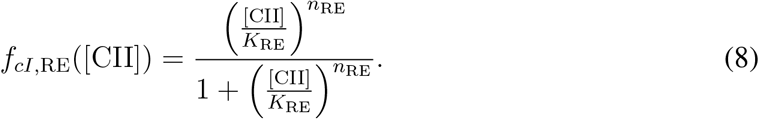

This function is used to define P_RE_ activity in the main text (**Figure 4A**).

CI activates its own expression from P_RM_ at low concentration, but represses P_RM_ at high concentration [14]. We assume competitive binding for Cro and CI at P_RM_, in line with previous models [1, 4, 15]. We extend this competitive binding logic to CI-activated and CI-repressed states, modeling each with distinct Hill activation or repression terms. Defining α_RM_ as the fold-change for transcription from P_RM_ in the CI-activated state, the single phage transcription regulatory function for *cI* expression from P_RM_ is

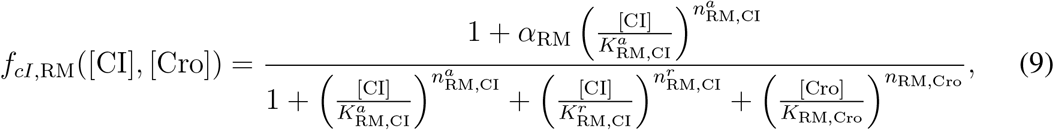

where α_RM_ is the fold-change for transcription from P_RM_ in the CI-activated state. The parameters for CI-repressed and CI-activated states are denoted with *r* and *a* superscripts for repression and activation respectively.

###### 2.1.2 *cro* Transcription

cro is transcribed from P_R_, and its transcription is repressed by itself and by CI. Employing the competitive binding assumption outlined in **Section 2.1.1**, the single phage transcription regulatory function is

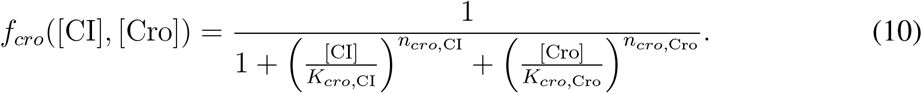

This function defines P_R_ activity in the main text (**Figure 6A**). To describe the strength of CI repression of *cro* transcription (main text, **Figure 4C**), we used the normalized weight of the CI-dependent term in Eq. 10, which we heuristically define as the probability of CI repressing cro:

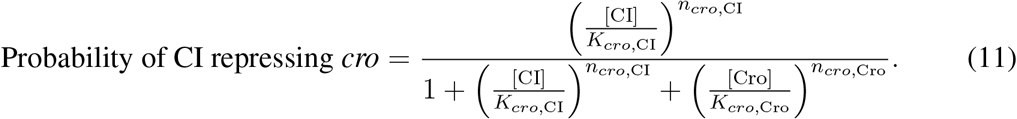

###### 2.1.3 *cII* Transcription

*cII* is also transcribed from P_R_, however its transcription requires readthrough of the terminator t_R1_ [10]. This readthrough is facilitated by the action of another lambda protein, N, that forms an antiterminator complex with host factors [10]. N is short-lived [10], and its transcription from promoter P_L_ is repressed by CI and Cro at concentrations similar to those that P_R_ is represssed at [14]. Based on this, the qualitative similarity in the *cro* and *cII* mRNA kinetics (main text, **Figure 2C**), and the observation that the termination efficiency at t_R1_ is relatively low (66% per [16]), we phenomenologically account for antitermination in our model by allowing the transcription regulatory function governing *cII* transcription to have parameters different from those describing *cro* transcription (Eq. 10). The single phage transcription regulatory function describing cII transcription is then

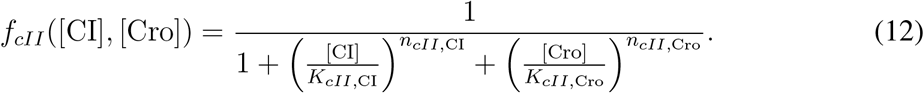

##### 2.2 CII Degradation

CII is actively degraded by the host protease FtsH, and has a half-life on the order of minutes

[10]. This degradation rate is reduced by the presence of another lambda protein, CIII, through an undetermined mechanism [17]. Transcription of *cIII* (from promoter P_L_) is repressed by both Cro and CI at concentrations similar to those that P_R_ is represssed at [14]. Like CII, CIII is also actively degraded by FtsH, and also has a half-life on the order of minutes [11]. Given the similarilty in transcriptional regulation and degradation of CII and CIII, we use CII concentration as a proxy for that of CIII. As a result the degradation regulatory function takes the following form:

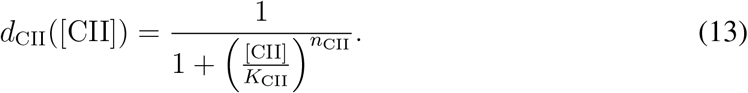

This functional form is motivated by a previous approach [12].

##### 2.3 Viral Replication

Viral copy number changes during infection as a result of viral replication [18, 19] which occurs through two primary modes: theta (also called circle-to-circle) and sigma (also called rolling circle) [11]. Lambda replication proteins O and P, which are transcribed from P_R_, are important for both modes of replication, and the dominant mode of replication switches from theta to sigma during infection [11]. Theta mode replication requires active transcription from P_R_ [10]. Repression of P_R_ by CI and Cro therefore prevents both theta mode of replication and transcription of O and P [10]. The molecular details of the different modes of replication and the switch from theta to sigma are still under investigation [11]. We follow the approach of [4,20] and coarse-grain replication into a single mode: exponential replication with a CI- and Cro-dependent rate given by the regulatory function

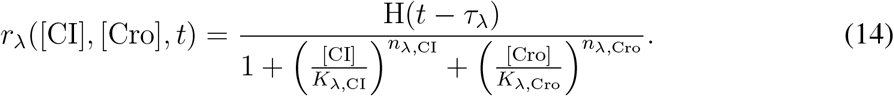

Notably, to account for the previously reported [19] delay in the onset of replication following infection (assumed to occur at *t* = 0), we have introduced the factor H(*t*-*τ*_λ_) with the Heaviside function (H(*t*) = 1 if t ≥ 0, 0 otherwise). Parameter *τ*_λ_ characterizes the duration of this delay. The functional form of Eq. 14 captures the repression of P_R_ by both CI and Cro. However, we allow the parameters used in this equation to differ from those in Eq. 10 to reflect the uncertainty in the molecular details of replication suppression.

To characterize the role of CI in repression of replication (main text, **Figure 4D**), we heuristically define the probability of CI repressing replication as the normalized weight of the CI-dependent term in the denominator of Eq. 14:

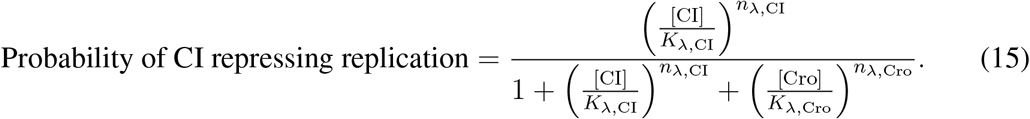

#### 3 Infection Simulation Methods

We numerically integrate the model ODEs (Eqs. 1–7 using MATLAB’s built-in ode15s solver, to account for possible stiffness due to nonlinear regulation terms and differences in time scales. For infection with *P*- phages, the replication rate in Eq. 7 is 0, and the ODE can be solved analytically. Defining initial cell volume as *V*_0_ (so that *V*(*t*) = V_0_exp(k_d_t)) and initial viral concentration as

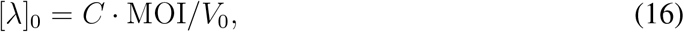

(where *C* is the conversion constant from volume to molar concentration), we express the viral concentration for *P*- infection as

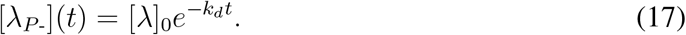

Model results in the main text are from simulations run for 60 minutes, which marks the end of the time series of *P*- mRNA measurements (**Figure 2C**, main text). All initial mRNA and protein concentrations are set to 0, and initial viral concentration (Eq. 16) is determined by the MOI and the initial cell volume. All concentrations are in units of nM.

To simulate the delayed infection scenario where infection by a second phage occurs *τ*_d_ minutes after initial infection (main text, **Figure 4**), the model ODEs in **Section 1** were integrated until *τ_d_* minutes after infection. Then we adjusted phage concentration by computing the corresponding change in viral concentration due to the addition of the second phage at the current volume, *V*(*τ*_d_). ODEs were then integrated from *τ_d_* to *t* = 60 min using the mRNA and protein concentrations at *τ_d_* and the updated viral concentration as the initial conditions.

#### 4 Parameter Fitting

Three data sets are used to fit the model: 1) *cI*, *cro*, and *cII* mRNA numbers from smFISH measurements during *P*- infection (described in the main text; see Experimental Methods for details), 2) measurements of viral copy number using qPCR during MOI = 1 *P*+ infection (also described in the main text; see Experimental Methods for details), and 3) previously published *cI* and *cII* mRNA numbers from smFISH measurements during infection with wild-type (*cI*+*cro*+*P*+), *cro*-, *cI*-, and *cro-P*- mutants (collectively denoted by subscript ‘Z’ in this text; see [19]).

All model parameters are fitted simultaneously, by minimizing the error between model output from simulations run for 60 minutes and experimental data, as described in the next section.

As we fit to population-averaged mRNA numbers, data points with only a single sample were ignored when fitting (see **Table S6**).

##### 4.1 Objective Function

Denoting model parameters by the vector ***θ***, and using hat notation for the model output (i.e., 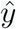), we fit the model to the data by minimizing the objective function

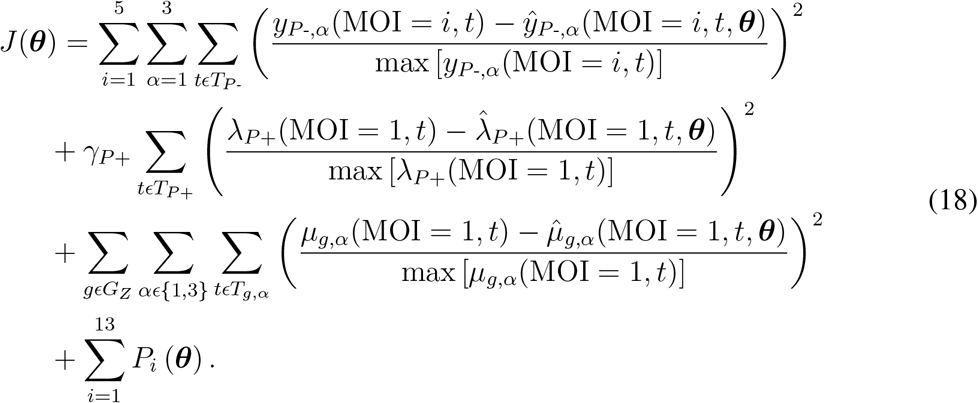

In Eq. 18, genotype is indicated in the subscript of fitted terms (e.g. *P*+ in *λ_P_*_+_), and the subscript *α ϵ* {1, 2, 3} denotes mRNA species (1 = *cI*, 2 = *cro*, and 3 = *cII*). The first term in Eq. 18 describes the error in fitting the *P*- data, the second term describes the error in fitting the *P*+ data, and the third term describes the error in fitting the previously published smFISH data. The last term describes penalties derived from qualitative relationships between parameters and observables described in the next section. To avoid problems with differences in scale in the residuals across data sets, we normalize each residual in the first three terms by the maximum of the relevant data subset.

While there may be differences between our smFISH experiments and those in [20] affecting total mRNA counts, we assumed that relative scaling in mRNA copy number between genotypes is preserved across experiments. To fit the previously published smFISH data, we thus compared relative differences between rescaled simulated and measured mRNA numbers, using maximum mRNA values in selected reference genotypes as normalization factors. These normalized terms are denoted *μ* in Eq. 18, and for a given genotype *g* and mRNA species *α* take the form

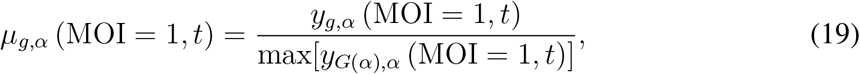

where *G*(*α*) denotes the reference genotype for mRNA species α. For *cII* mRNA measurements, we used the *P*- mutant as the reference genotype. For *cI* mRNA measurements, we used the *cro-P*- mutant, as *cI* mRNA during *P*- infection was not measured in [20].

The number of data points in our *P*+ replication data is significantly smaller (7 data points) than that in either the *P*- smFISH measurements (150 data points) or the previously published smFISH data (50 data points). Therefore, to ensure simultaneous fits to all data captured trends in the replication data, we assigned an additional weight (*γ*_*P*+_ = 8) to the *P*+ error term. The value of this weight approximately corresponds to the ratio in the number of points between the previously published smFISH and *P*+ data sets.

##### 4.2 Penalties

While most parameters in the model have never been directly measured, qualitative relationships between some parameters can be gleaned from previous experimental measurements, resulting in inequality constraints. Such constraints can also be deduced for the kinetics of some observables, such as viral copy number. Because particle swarm optimization (described in **Section 4.3**) does not require a differentiable objective function, we directly penalized violations of inequality constraints using a combination of simple Heaviside functions and quadratic terms (motivated by exact penalization in direct search methods—see [21, 22]). To avoid scaling issues, we normalized the quadratic penalties by the larger value in the violated inequality. We also included ‘count’ penalties [23] to ensure penalty terms have a non-negligible weight in the objective function even when the violation of the inequality is small. For a given argument pair, *x_j_* (***θ***) and *x_k_* (**θ**), a penalty term based on the strict inequality constraint *x_k_* (***θ***) < *x_j_* (***θ***) has the form

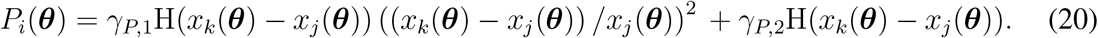

Here the first term corresponds to the quadratic penalty for the constraint, the second term is the count penalty, and H(x) is the Heaviside function.

The penalty weights (*γ_P,1_* and *γ_P,2_*) were both fixed at 0.1, a value empirically found to result in fits that satisfy constraints without compromising fit quality. A full list of parameter constraints is given in **Theoretical Methods Table 1 (Section 7**).

##### 4.3 Particle Swarm Optimization

We minimized the objective function in Eq. 18 using MATLAB’s built-in algorithm for particle swarm optimization (PSO). PSO is a global optimization algorithm [24] that performs well for a wide range of optimization problems [25–27]. Optimization runs were carried out using randomized initial positions in parameter space, and the fitting code was parallelized to run on the NOTS high performance computing cluster at Rice University. Internal parameters of the PSO algorithm that were modified from their default values in MATLAB are listed in **Theoretical Methods Table 2 (Section 7**).

We use the top 25% of fits (as measured by the value of the objective function) obtained from end-points of multiple separate PSO runs for our ensemble of parameter sets.

#### 5 Decision Thresholds

Ranges for CI and Cro thresholds for the lysis-lysogeny decision were obtained using previously published observations on the frequency of each fate.

For *P*+ infection, lysis is the dominant outcome during at MOI = 1 [28]. However, lysis has not been observed with *P*- phages, either in bulk [29] or in single-cell resolution experiments over a range of MOI [19]. Based on these observations, we used our model simulations to constrain possible Cro thresholds for lysis:

- Maximum lytic threshold: the maximum Cro concentration reached in simulations of *P*+ infection at MOI = 1,
- Minimum lytic threshold: the maximum Cro concentration reaching in simulations of *P*- infection at MOI = 5.

To constrain the lysogenic threshold, we first defined the MOI at which the decision switches from lysis to lysogeny during *P*+ infection. As discussed in the main text, we interpret the observed separation of trajectories at MOI = 1 and MOI = 2 (main text, **Figure 3B**) as an indication of a switch in decision outcome. Therefore, we assume that the lysogenic threshold for CI is reached at MOI= 2 and not reached at MOI= 1 Based on this interpretation, we obtain the following constraints for the lysogenic threshold:

- Maximum lysogenic threshold: the maximum CI concentration reached in simulations of *P*+ infection at MOI = 2.
- Minimum lysogenic threshold: the maximum CI concentration reached in simulations of *P*+ infection at MOI = 1.

Notably, this identification of a transition in infection outcome from MOI = 1 to MOI = 2 is consistent with a previous simple model of Kourilsky et al. [30], who also found that the dominant outcome switches to lysogeny at MOI = 2 during *P*+ infection. More recent lambda models have also assumed the lysis-to-lysogeny transition occurs at MOI = 2 based on this previous simple model [1, 12, 31].

To prevent cases where neither threshold is reached during *P*+ infection for 1 < MOI < 2 when scanning MOI as a continuous variable for the fate diagram (main text, **Figure 6E**), we scanned over MOI in this range, decreasing the maximum values of lytic or lysogenic thresholds further until a threshold is always crossed.

In principle, any decision threshold within the defined bounds can lead to the prediction of cell fate that will satisfy the above-mentioned constraints. For simplicity and to achieve the maximally robust predictions, we set the lytic and lysogenic thresholds to the midpoints of their respective threshold ranges for all figures shown in the main text. We note that while changes in the values of the thresholds may affect the size and exact boundaries of fate regions in **Figure 6E** (main text), the topology of the fate diagram is conserved. For example, changes in the value of the lysogenic threshold affect the value of MOI at which the infection switches from failed infection to lysogeny during *P*- infection from 3 to 4. On the other hand, changes in the lytic threshold affect the MOI at which the mixed outcome regime occurs (grey region in **Figure 6E**, main text).

### 6 Modeling Bulk Lysogenization

The fraction of cells undergoing lysogeny as a function of the population-averaged MOI was measured as described in the **Experimental Methods (Section 4.1)**. To estimate MOI*, the single-cell MOI at which the transition to lysogeny occurs, we followed the approach of [28, 30, 32]. First, we assume that phage-cell encounters in the infection mixture follow Poisson statistics. Therefore, the probability that a cell is infected by *n* phages, given an average MOI of *M*, is:

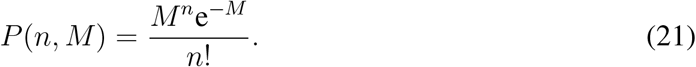

We next assume that coinfection by MOI* phages or more results in lysogeny, while coinfection by fewer than MOI* does not. In other words, the probability *Q* that a cell is lysogenized at single-cell MOI = *n* follows:

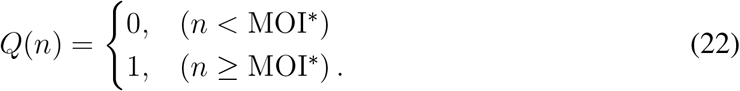

The model-predicted lysogenization frequency 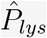, given an average MOI of *M*, is then found by summation over all possible values of *n*:

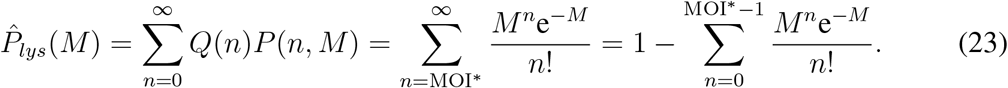

To evaluate MOI*, we fitted Eq. 23 to the experimentally measured lysogenization frequencies *P_lys_*(*M*) by minimizing the objective function:

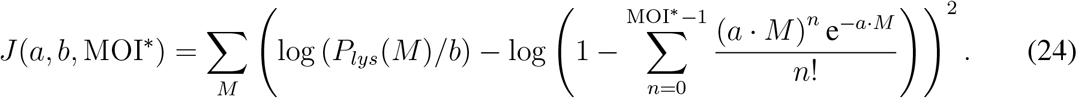

Here, *a* and *b* are fitted normalization factors accounting for experiment-to-experiment errors in measuring the absolute numbers of phages and bacteria, and MOI* is allowed to take the values 1, 2, and 3. Fitting was performed in logarithmic space, since the serial dilutions used to scan bulk MOI result in data spanning multiple orders of magnitude. **Figure 6D** depicts the best fit for each dataset, with the experimental data rescaled by the fitting parameters, (i.e., *1/b* · *P_lys_*(*M*) vs. *a* · *M*).

### 7 Theoretical Methods Tables

**Theoretical Methods Table 1:** Description of constraints used in fitting

| Constraint | Description | Reference |
| --- | --- | --- |
| $k_{tx}^{1, \text{RM}} \alpha_{\text{RM}} < k_{tx}^{1, \text{RE}}$ | $P_{\text{RE}}$ has a higher maximum transcription rate than $P_{\text{RM}}$ | Herskowitz and Hagen, Annu. Rev. Genet., 1980 |
| $k_{tx}^{1, \text{RM}} \alpha_{\text{RM}} < k_{tx}^2$ | $P_{\text{R}}$ has a higher maximum transcription rate than $P_{\text{RM}}$ | Dodd et al., Genes and Dev., 2004 |
| $k_{tx}^{1, \text{RE}} < k_{tx}^2$ | $P_{\text{R}}$ has a higher maximum transcription rate than $P_{\text{RE}}$ | Palmer et al., Mol. Cell, 2009 |
| $k_{tr}^1 < k_{tr}^2$ | CI is translated at a slower rate than Cro | Liu et al., PNAS, 2013 |
| $k_{tr}^1 < k_{tr}^3$ | CI is translated at a slower rate than CII | Liu et al., PNAS, 2013 |
| $K_{cro, \text{CI}} < K_{\text{RM}, \text{CI}}^a$ | CI represses *cro* transcription before activating its own transcription | Ptashne, The Genetic Switch, 1986 |
| $K_{\text{RM}, \text{CI}}^a < K_{\text{RM}, \text{CI}}^r$ | CI activates its own transcription at lower concentration than that at which it represses its own transcription | Ptashne, The Genetic Switch, 1986 |
| $K_{cII, \text{CI}} < K_{\text{RM}, \text{CI}}^a$ | CI represses *cII* transcription before activating its own transcription | Ptashne, The Genetic Switch, 1986 |
| $K_{\text{RM}, \text{Cro}} < K_{cro, \text{Cro}}$ | Cro represses $P_{\text{RM}}$ before repressing its own transcription | Ptashne, The Genetic Switch, 1986 |
| $K_{\text{RM}, \text{Cro}} < K_{cII, \text{Cro}}$ | Cro represses $P_{\text{RM}}$ before repressing *cII* transcription | Ptashne, The Genetic Switch, 1986 |
| $K_{cro, \text{CI}} \leq K_{\lambda, \text{CI}}$ | CI suppression of replication occurs at or above the concentration threshold for repression of $P_{\text{R}}$ | Oppenheim, Annu. Rev. Genet., 2005 |
| $K_{cro, \text{Cro}} \leq K_{\lambda, \text{Cro}}$ | Cro suppression of replication occurs at or above the concentration threshold for repression of $P_{\text{R}}$ | Oppenheim, Annu. Rev. Genet., 2005 |
| $\frac{\lambda_{P+}(\text{MOI}=1, t=40 \text{ min})}{\lambda_{cro-}(\text{MOI}=1, t=40 \text{ min})} > 5$ | At least 5-fold more viruses are produced by 40 min during *P+* infection vs. *cro-* infection | Shao et al., iScience, 2018 |

**Theoretical Methods Table 2:**
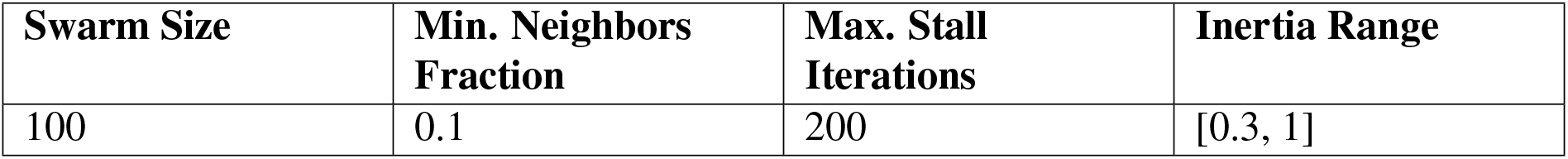
Internal particle swarm optimization parameters changed from MATLAB’s built-in defaults.

| **Swarm Size** | **Min. Neighbors Fraction** | **Max. Stall Iterations** | **Inertia Range** |
| --- | --- | --- | --- |
| 100 | 0.1 | 200 | [0.3, 1] |

**Theoretical Methods Table 3:** Fitted parameters used for main text figures (with *V_0_* = 1*μ*m^3^).

| Parameter | Description | Value (main text) |
| --- | --- | --- |
| $k_{tx}^{1,RM}$ | Basal *cI* transcription rate from $P_{RM}$ | $0.205 \text{ min}^{-1}$ |
| $\alpha_{RM}$ | Fold-change in $P_{RM}$ activity | 5.00 |
| $k_{tx}^{1,RE}$ | *cI* transcription rate from $P_{RE}$ | $1.04 \text{ min}^{-1}$ |
| $k_{tr}^1$ | *cI* mRNA translation rate | $0.887 \text{ min}^{-1}$ |
| $k_{tx}^2$ | *cro* transcription rate | $2.54 \text{ min}^{-1}$ |
| $k_{tr}^2$ | *cro* mRNA translation rate | $0.919 \text{ min}^{-1}$ |
| $k_{tx}^3$ | *cII* transcription rate | $5.29 \text{ min}^{-1}$ |
| $k_{tr}^3$ | *cII* mRNA translation rate | $1.10 \text{ min}^{-1}$ |
| $k_\lambda$ | Viral replication rate | $0.131 \text{ min}^{-1}$ |
| $k_d$ | Dilution rate | $0.250 \text{ min}^{-1}$ |
| $k_m^1$ | *cI* mRNA degradation rate | $0.100 \text{ min}^{-1}$ |
| $k_P^1$ | CI degradation rate | $3.22 \times 10^{-9} \text{ min}^{-1}$ |
| $k_m^2$ | *cro* mRNA degradation rate | $0.158 \text{ min}^{-1}$ |
| $k_P^2$ | Cro degradation rate | $1.33 \times 10^{-9} \text{ min}^{-1}$ |
| $k_m^3$ | *cII* mRNA degradation rate | $0.100 \text{ min}^{-1}$ |
| $k_P^3$ | CII degradation rate | $0.369 \text{ min}^{-1}$ |
| $n_{RM,CI}^a$ | Hill function sensitivity parameter for CI activation of $P_{RM}$ | 2.00 |
| $n_{RM,CI}^r$ | Hill function sensitivity parameter for CI repression of $P_{RM}$ | 6.00 |
| $n_{RM,Cro}$ | Hill function sensitivity parameter for Cro repression of $P_{RM}$ | 3.00 |
| $n_{RE}$ | Hill function sensitivity parameter for CII activation of $P_{RE}$ | 5.00 |
| $n_{cro,CI}$ | Hill function sensitivity parameter for CI repression of *cro* transcription | 2.55 |
| $n_{cro,Cro}$ | Hill function sensitivity parameter for Cro repression of *cro* transcription | 2.22 |
| $n_{cII,CI}$ | Hill function sensitivity parameter for CI repression of *cII* transcription | 3.95 |
| $n_{cII,Cro}$ | Hill function sensitivity parameter for Cro repression of *cII* transcription | 3.00 |
| $n_{\lambda,CI}$ | Hill function sensitivity parameter for CI repression of viral replication | 6.00 |
| $n_{\lambda,Cro}$ | Hill function sensitivity parameter for Cro repression of viral replication | 6.00 |
| $n_{CII}$ | Hill function sensitivity parameter for suppression of CII degradation | 1.00 |
| $K_{RM,CI}^a$ | Hill function threshold parameter for CI activation of $P_{RM}$ | 108 nM |
| $K_{RM,CI}^r$ | Hill function threshold parameter for CI repression of $P_{RM}$ | 528 nM |
| $K_{RM,Cro}$ | Hill function threshold parameter for Cro-repression of $P_{RM}$ | 20.4 nM |
| $K_{RE}$ | Hill function threshold parameter for CII activation of $P_{RE}$ | 78.1 nM |
| $K_{cro,CI}$ | Hill function threshold parameter for CI repression of *cro* transcription | 64.1 nM |
| $K_{cro,Cro}$ | Hill function threshold parameter for Cro repression of *cro* transcription | 94.2 nM |
| $K_{cII,CI}$ | Hill function threshold parameter for CI repression of *cII* transcription | 64.1 nM |
| $K_{cII,Cro}$ | Hill function threshold parameter for Cro repression of *cII* transcription | 94.2 nM |
| $K_{\lambda,CI}$ | Hill function threshold parameter for CI repression of viral replication | 241 nM |
| $K_{\lambda,Cro}$ | Hill function threshold parameter for Cro repression of viral replication | 500 nM |
| $K_{CII}$ | Hill function threshold parameter for CII degradation suppression | 241 nM |
| $\tau_\lambda$ | Time offset for onset of replication | $7.50 \text{ min}^{-1}$ |

## Supplementary Figures

**Figure S1.**
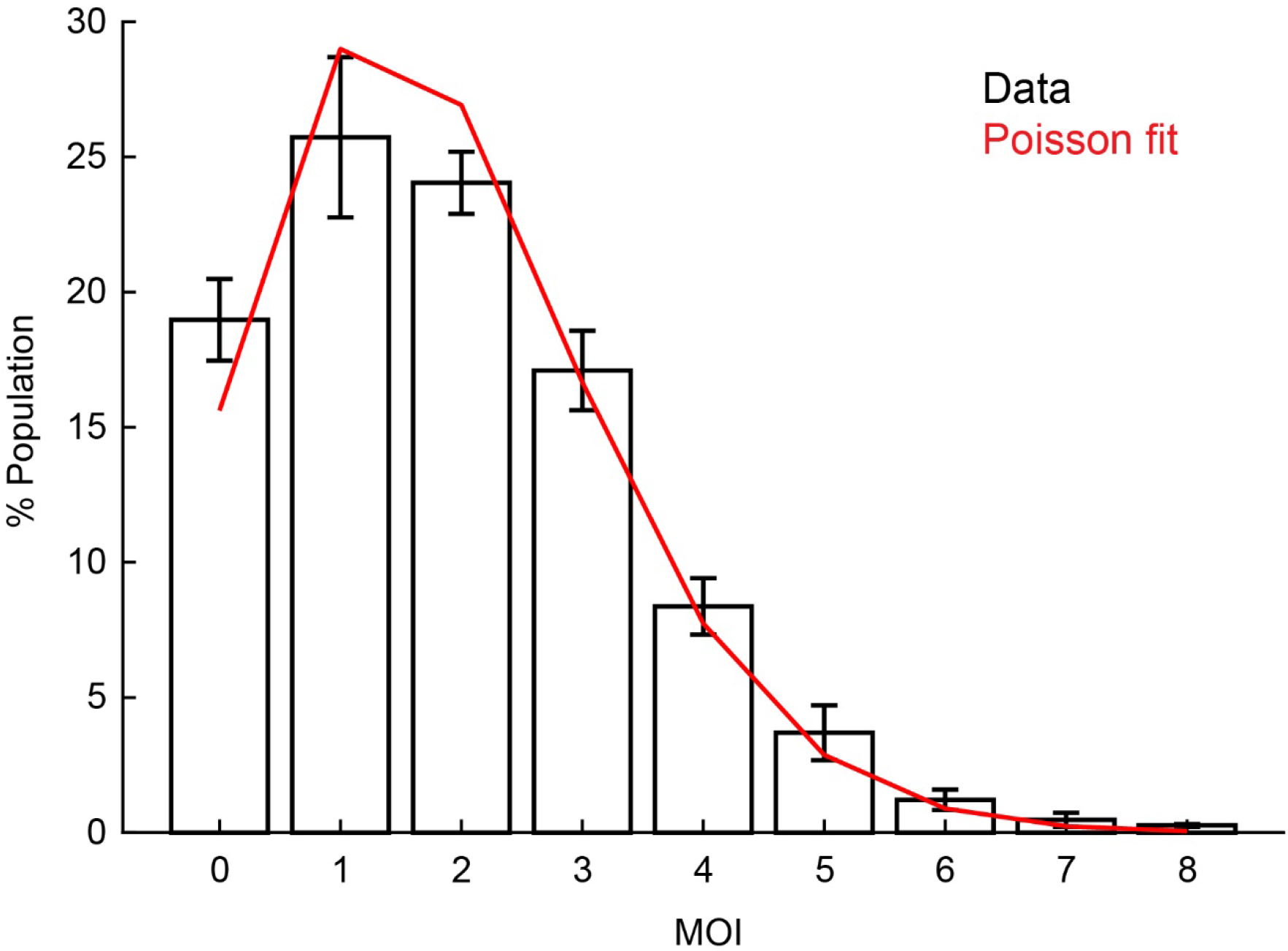
The distribution of single-cell MOI within a population. The distribution of single-cell MOI following infection by *λ cI*857 *Pam*80 *P1parS*. The infection procedure is described in **Experimental Methods Section 4.2**, and the identification of single-cell MOI in **Experimental Methods Section 8.2**. Markers and error bars indicate mean ± SEM from samples taken at 1, 2 and 5 minutes following infection (see **Table S6** for detailed sample sizes). The mean MOI calculated from these samples was 1.9 ± 0.5 (mean ± SEM). Red line: Poisson distribution of the same mean, reflecting the assumption of random encounters between phages and bacteria (Kourilsky, P., Mol Gen Genet, 1973).

**Figure S2.**
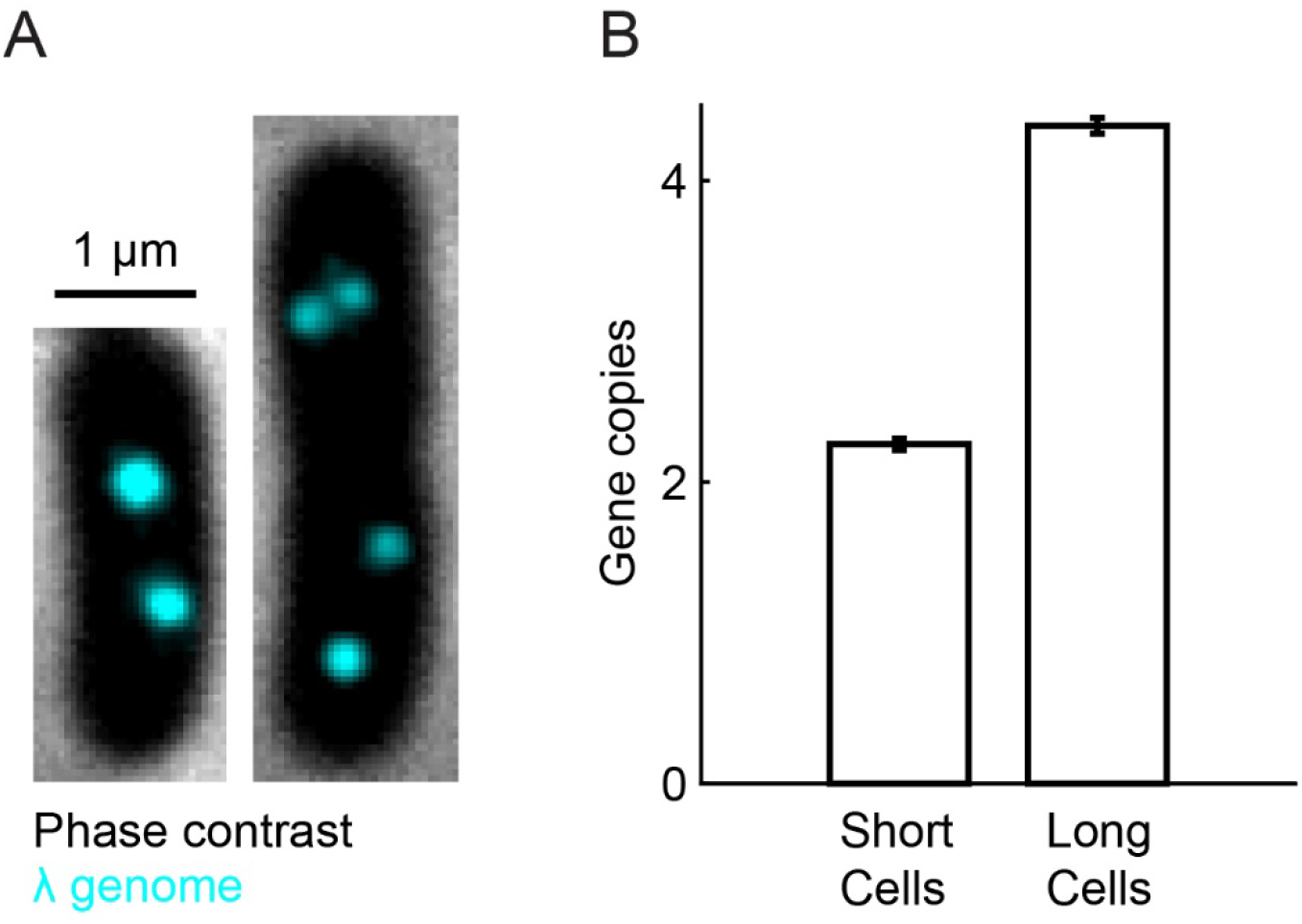
The estimated number of lambda prophage copies in lysogenic cells. **(A)** Lysogenic strain MG1655 carrying prophage *λ cI*857 *ind^-^ P1parS* and plasmid pALA3047 (expressing CFP-ParB). Cells were grown at 30°C in LBMM supplemented with 10 μM IPTG. Individual prophages are labeled using CFP-ParB. The imaging procedure is described in **Experimental Methods Section 6**. **(B)** Newborn lysogenic cells (“short cells”, defined as the 5-20 percentiles of cell lengths, N = 3-50) contain two lambda genome copies as expected (Bremer, H. and Churchward, G., J Theor Biol, 1977), while cells about to divide (“long cells”, the 80-95 percentiles of cell length, N = 349) contain four copies. Error bars indicate SEM.

**Figure S3.**
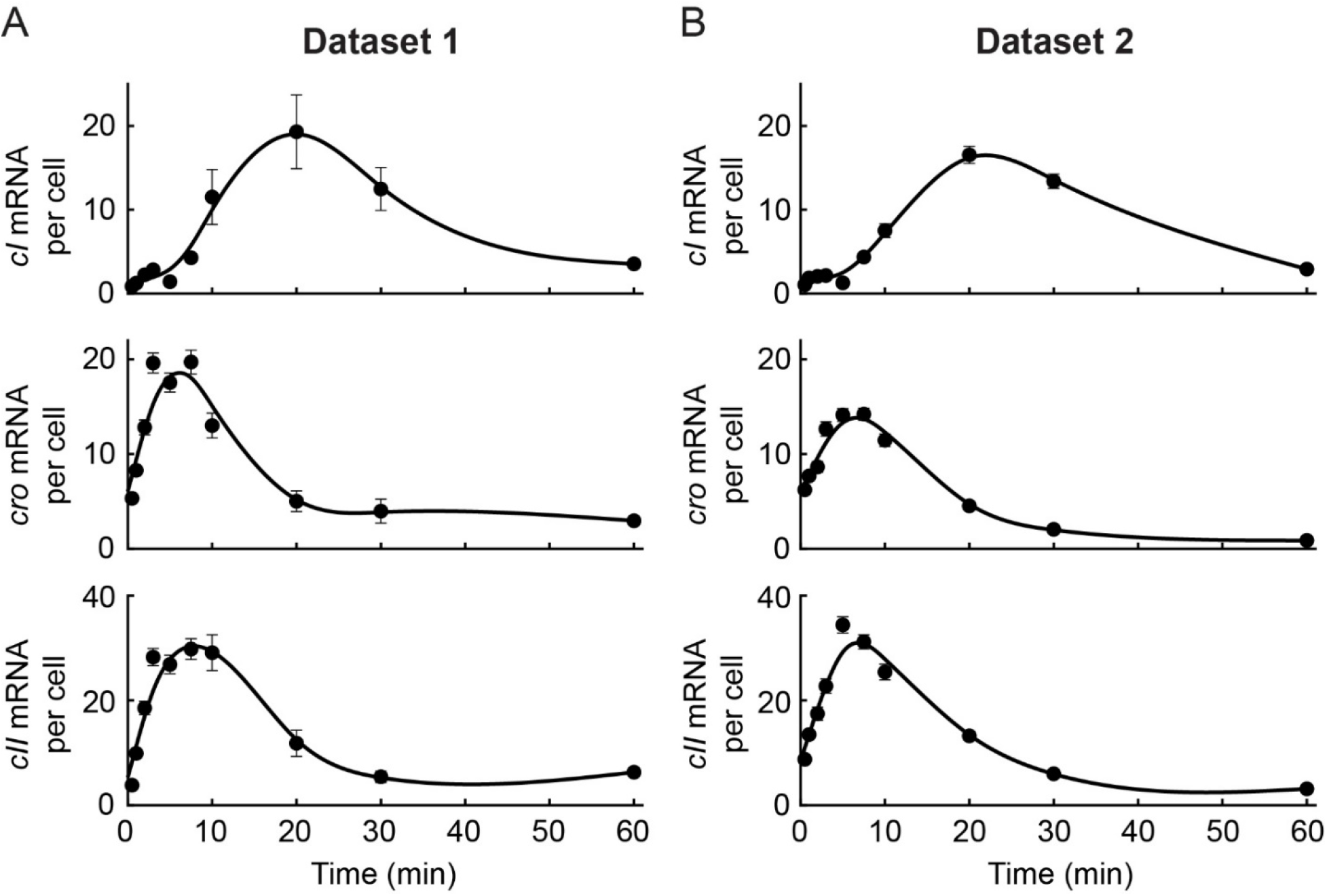
The transcription kinetics of *cI, cro* and *cII* across biological replicates. The numbers of *cI, cro*, and *cII* mRNA per cell (mean ± SEM), at different times following infection by *λ cI*857 *Pam*80 *P1parS*. The infection procedure is described in **Experimental Methods Section 4.2**, and the mRNA quantification in **Experimental Methods Section 8.1**. **(A)** Results from dataset 1 (see **Table S6** for detailed sample sizes). The mean MOI was 2.1 ± 0.3 (SEM). **(B)** Results from dataset 2 (see **Table S6** for detailed sample sizes). The mean MOI was 2.1 ± 0.2 (SEM). Solid lines are splines, used to guide the eye.

**Figure S4.**
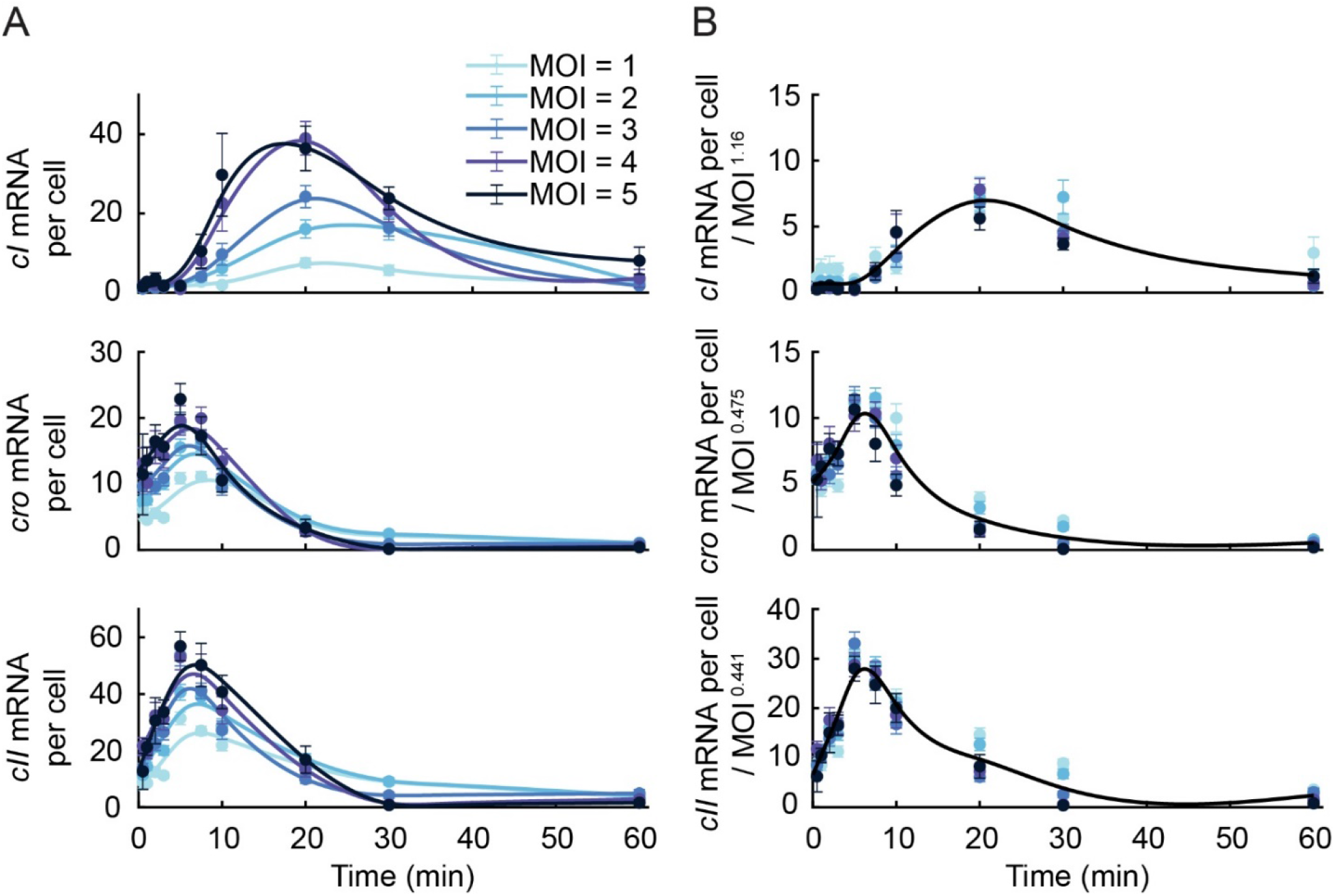
MOI scaling of *cI*, *cro* and *cII* trajectories. **(A)** The numbers of *cI*, *cro*, and *cII* mRNA per cell (mean ± SEM), at different times following infection by λ *cI*857 *Pam*80 *PlparS*, at single-cell MOI of 1–5. The infection procedure is described in **Experimental Methods Section 4.2**, the mRNA quantification and single-cell MOI identification in **Experimental Methods Sections 8.1** and **8.2**, respectively. Solid lines indicate splines, used to guide the eye. **(B)** The values in panel A, scaled by MOI^ε^, with ε equal 1.16, 0.475, 0.441 for *cI*, *cro* and *cII*, respectively. The optimal value of ε for each gene was obtained by minimizing the sum of squared deviation between mRNA values at different MOI at all time points. Solid lines indicate a single spline over all scaled mRNA numbers.

**Figure S5.**
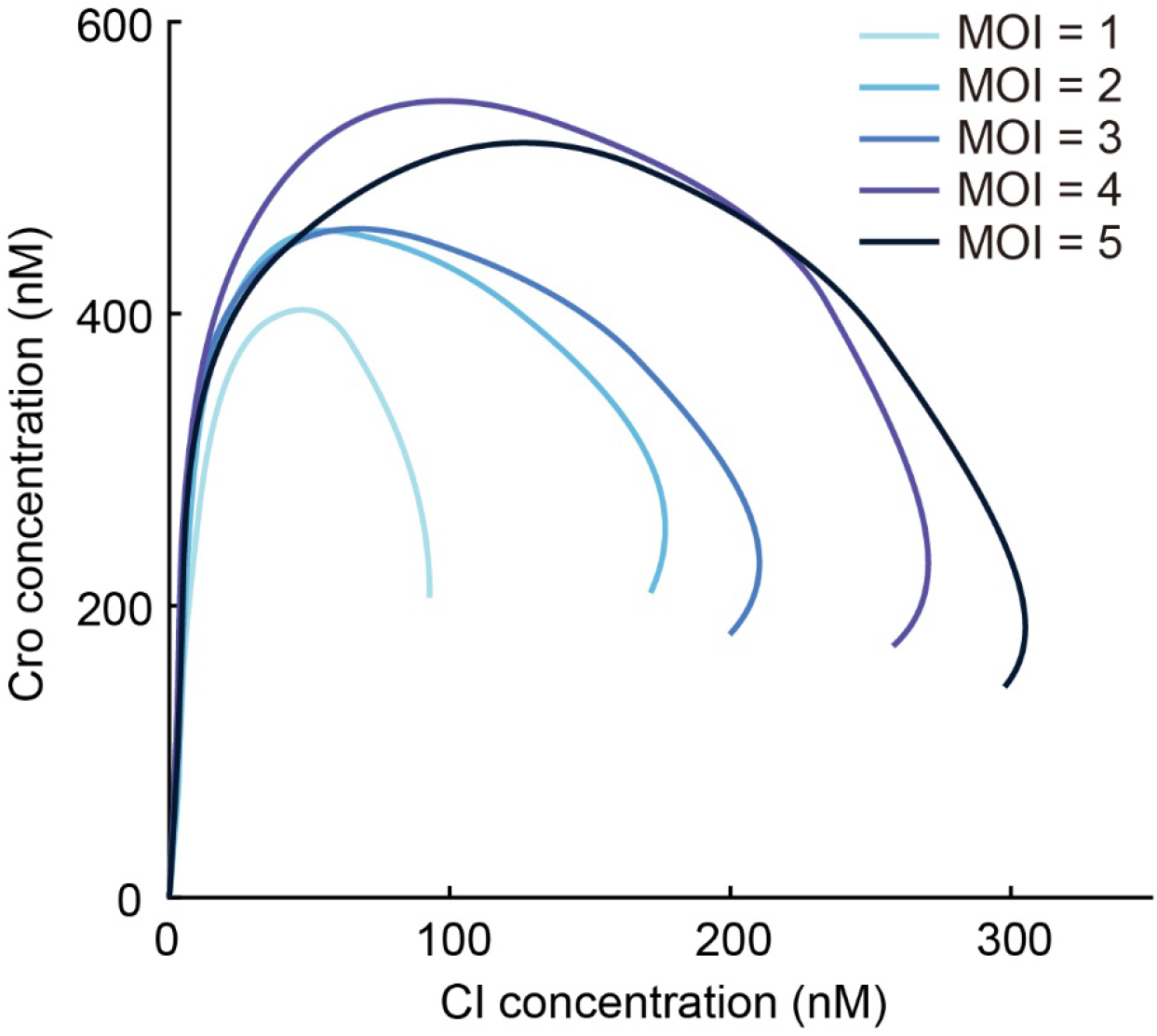
Estimated CI-Cro trajectories following infection. The estimated trajectories in the plane of Cro and CI concentration, during the first 60 minutes following infection by *λ cI*857 *Pam*80 *P1parS*, at single-cell MOI of 1–5. The infection procedure is described in **Experimental Methods Section 4.2**, the mRNA quantification and single-cell MOI identification in **Experimental Methods Sections 8.1** and **8.2**, respectively. We first calculated the concentration of *cI* and *cro* mRNA, [mRNA], by dividing the number of molecules in each cell by the cell volume (approximated as a spherocylinder, with dimensions obtained from the automated segmentation, see **Experimental Methods Section 7.1**). [mRNA] was then used to estimate the protein concentration of the corresponding species using the relation: d[protein]/dt = translation rate × [mRNA] – decay rate × [protein], with the rates of translation and decay taken from the literature (Zong, C., et al., Mol Syst Biol, 2010; Reinitz, J. and Vaisnys J.R., J Theor Biol, 1990).

**Figure S6.**
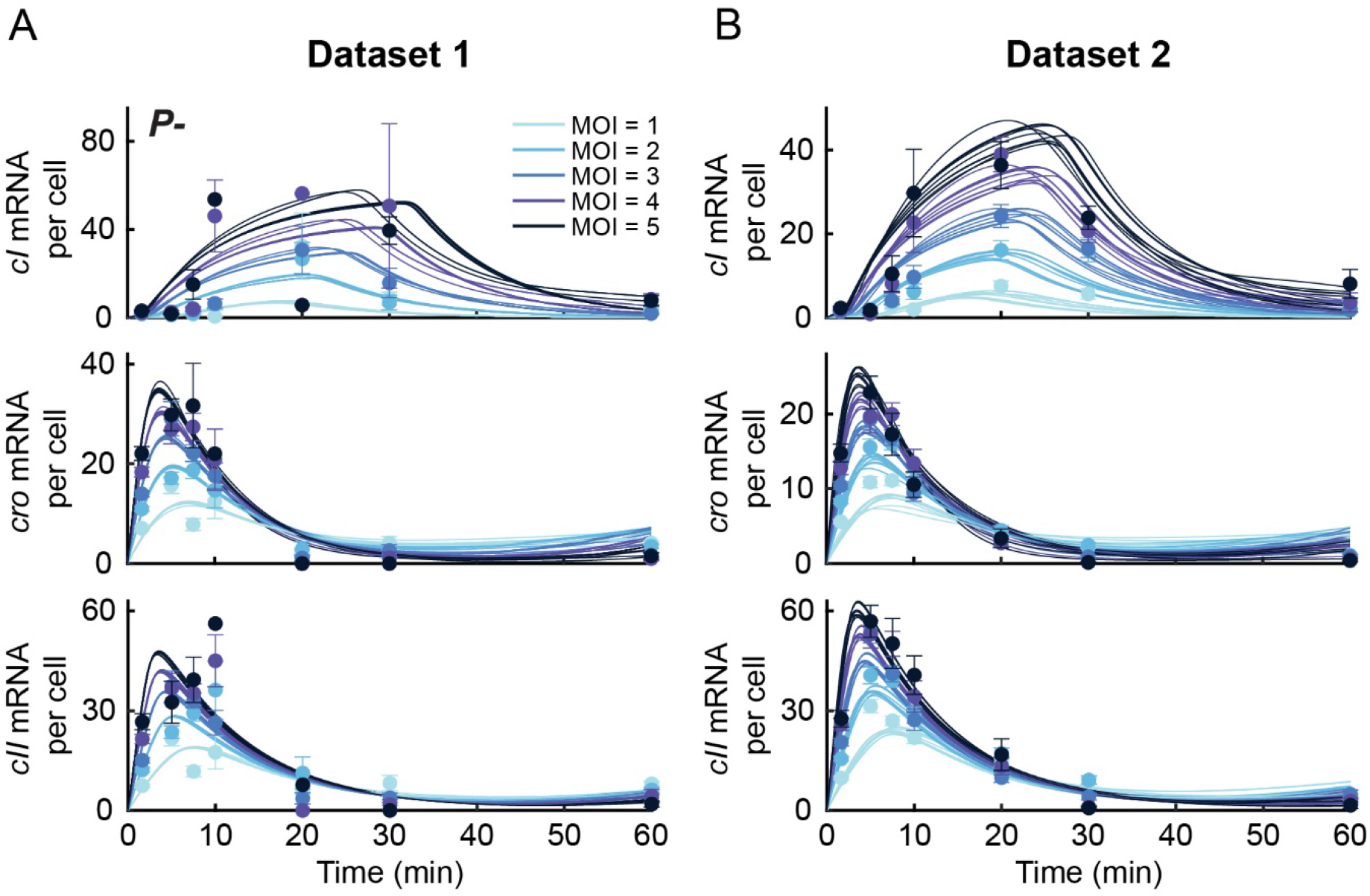
Model trajectories from ensembles of fits capture mRNA kinetics in biological replicates of *P*- infection. The numbers of *cI*, *cro*, and *cII* mRNA per cell, at different times following infection at MOI = 1–5 by *P*- phage (*λ cI*857 *Pam*80 *P1parS*; see also **Figure 2C**, main text). **(A)** Results from dataset 1. **(B)** Results from dataset 2. Markers and error bars indicate experimental mean ± SEM of each sample. Solid lines indicate ensembles of model fits that yield consistent predictions, obtained from minimizing the objective function described in **Theoretical Methods Section 4**. See **Table S6** for samples sizes.

**Figure S7.**
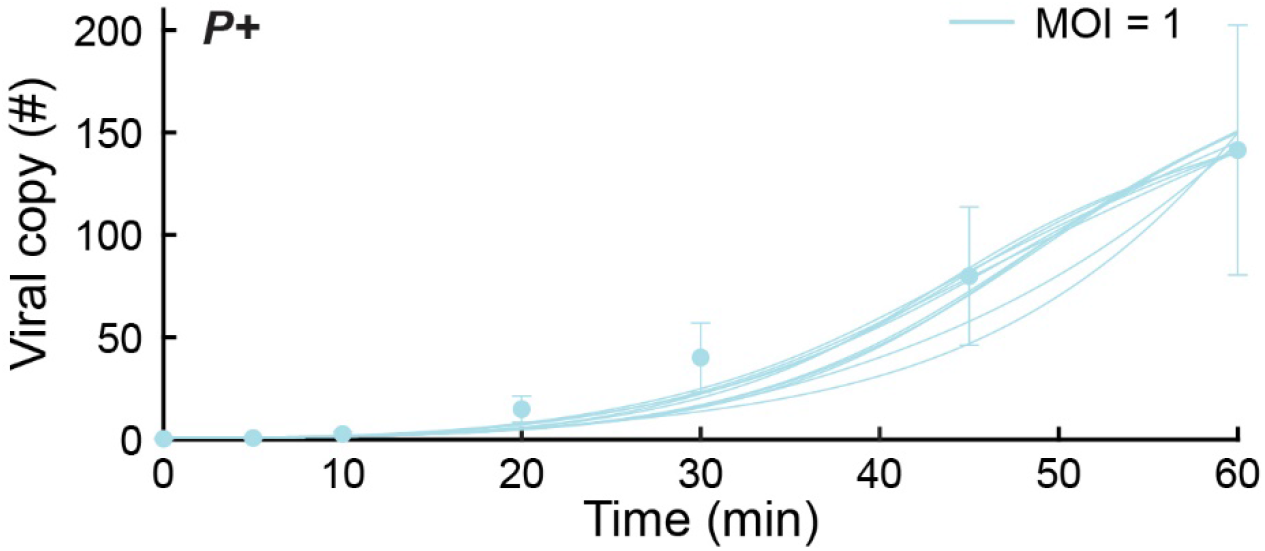
Model trajectories from an ensemble of fits capture the dynamics of viral copy number during *P*+ infection. Viral copy number, measured using qPCR, following infection at MOI = 1 by *P*+ phage (*λ cI*857 *Sam*7; see also **Figure 2D**, main text). Markers and error bars indicate experimental mean ± standard deviation due to qPCR calibration uncertainty. Solid lines indicate an ensemble of model fits obtained from minimizing the objective function described in **Theoretical Methods Section 4**, using the *P*- mRNA measurements from dataset 2 (**Figure S6**).

**Figure S8.**
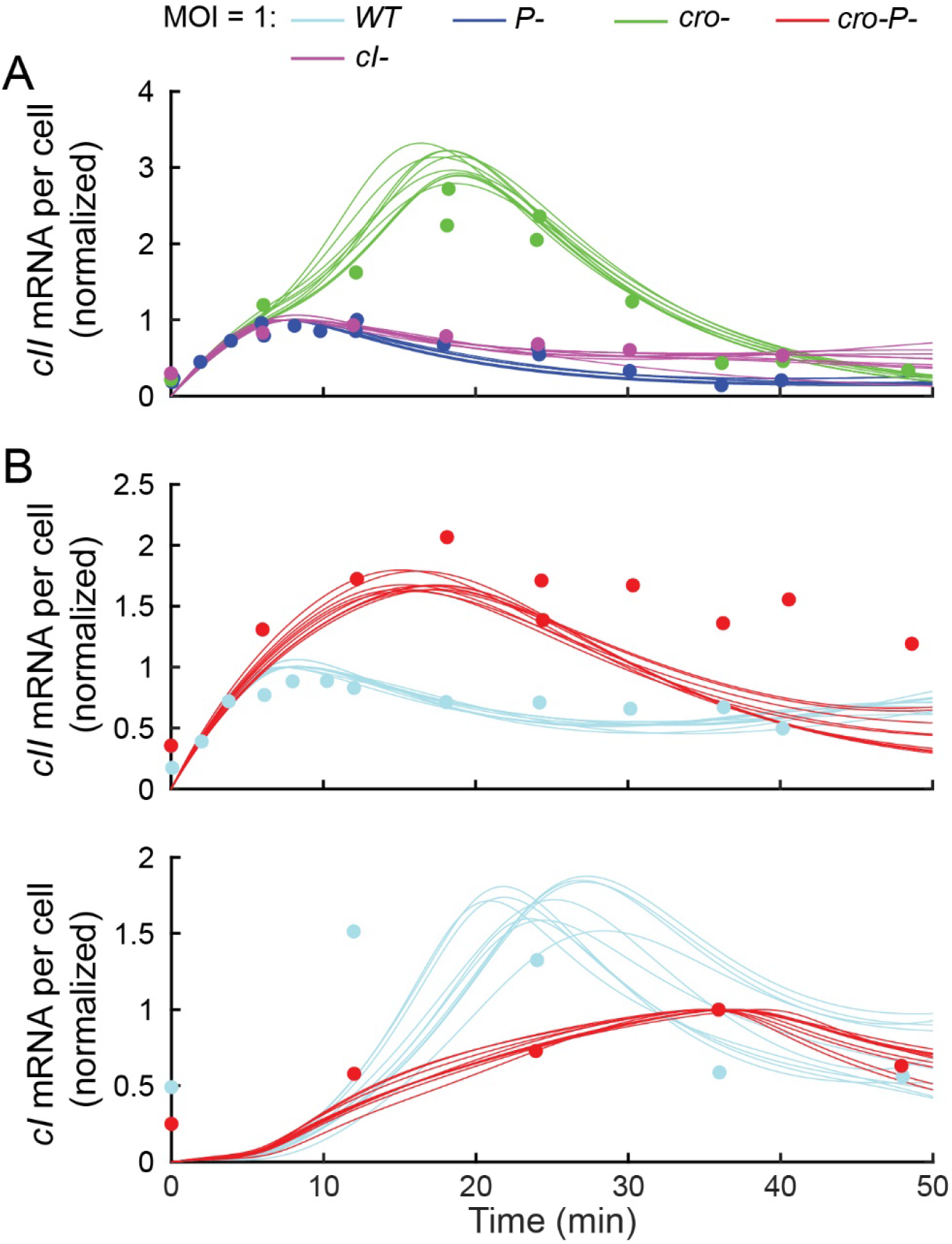
Model trajectories from an ensemble of fits capture *cI* and *cII* mRNA kinetics during infections with various lambda genotypes. The numbers of *cII* mRNA (normalized by maximum *cII* mRNA count during *P*- infection) and *cI* mRNA (normalized by maximum *cI* mRNA count during *cro-P*- infection) per cell at different times following infection at MOI = 1 with various lambda genotypes (data from Shao, Q. et al., iScience, 2018). **(A)** *cII* mRNA following infection with *P*-, *cI*-, and *cro*- phages. **(B)** *cII* (top) and *cI* (bottom) mRNA following infection with WT (*cI*+*cro*+*P*+) and *cro-P*- phages. Solid lines indicate an ensemble of model fits obtained from minimizing the objective function described in **Theoretical Methods Section 4**, using the *P*- mRNA measurements from dataset 2 (**Figure S6**).

**Figure S9.**
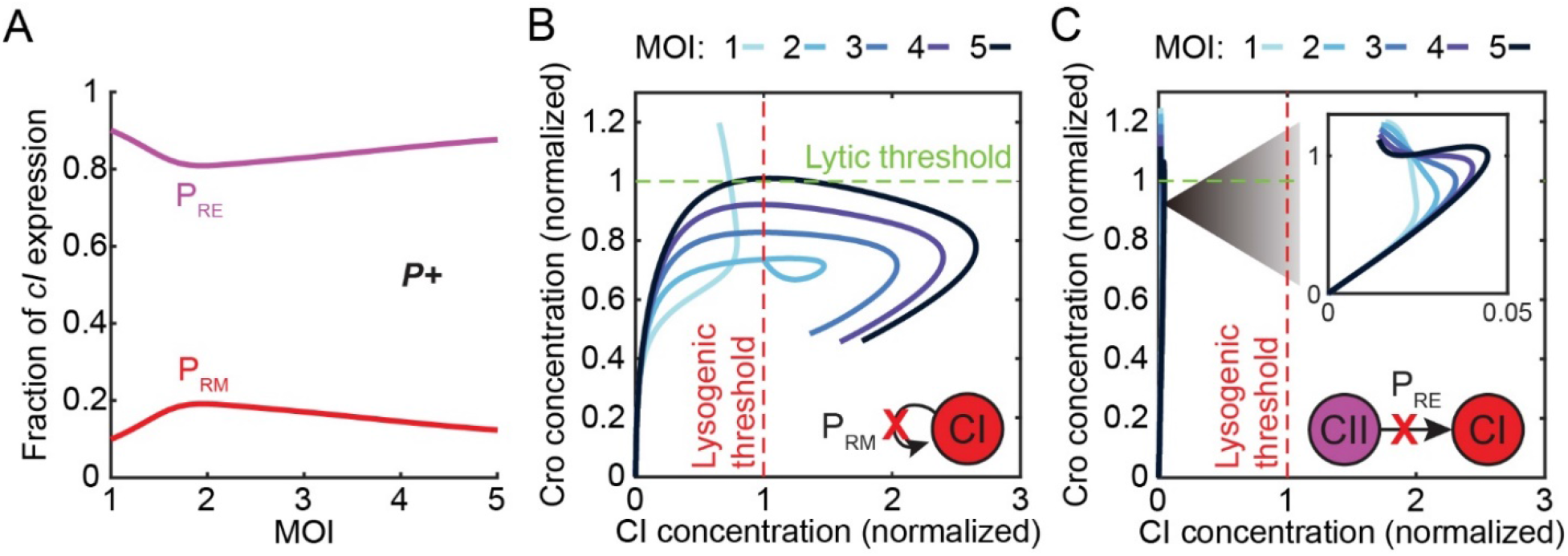
CII-activated *cI* expression from P_RE_ is required for a lysis-to-lysogeny transition, while *cI* autoactivation is not. **(A)** The model-predicted fraction of total *cI* mRNA expressed from P_RE_ (purple) and P_RM_ (red) during the first 60 minutes of infection with *P*+ phages over a range of MOI. The majority of *cI* expression comes from P_RE_ for all MOI simulated. **(B)** Model-predicted trajectories, in the plane of Cro and CI concentrations, during the first 60 minutes following infection by phages in which *cI* autoactivation of P_RM_ has been removed. A lysis-to-lysogeny transition is achieved even in the absence of CI-activated transcription from P_RM_. **(C)** Same as panel B, for the case of infection by a phage in which CII activation of P_RE_ has been removed. In the absence of *cI* transcription from P_RE_, a transition to lysogeny is not observed. Protein concentrations were normalized by the lytic and lysogenic thresholds.

**Figure S10.**
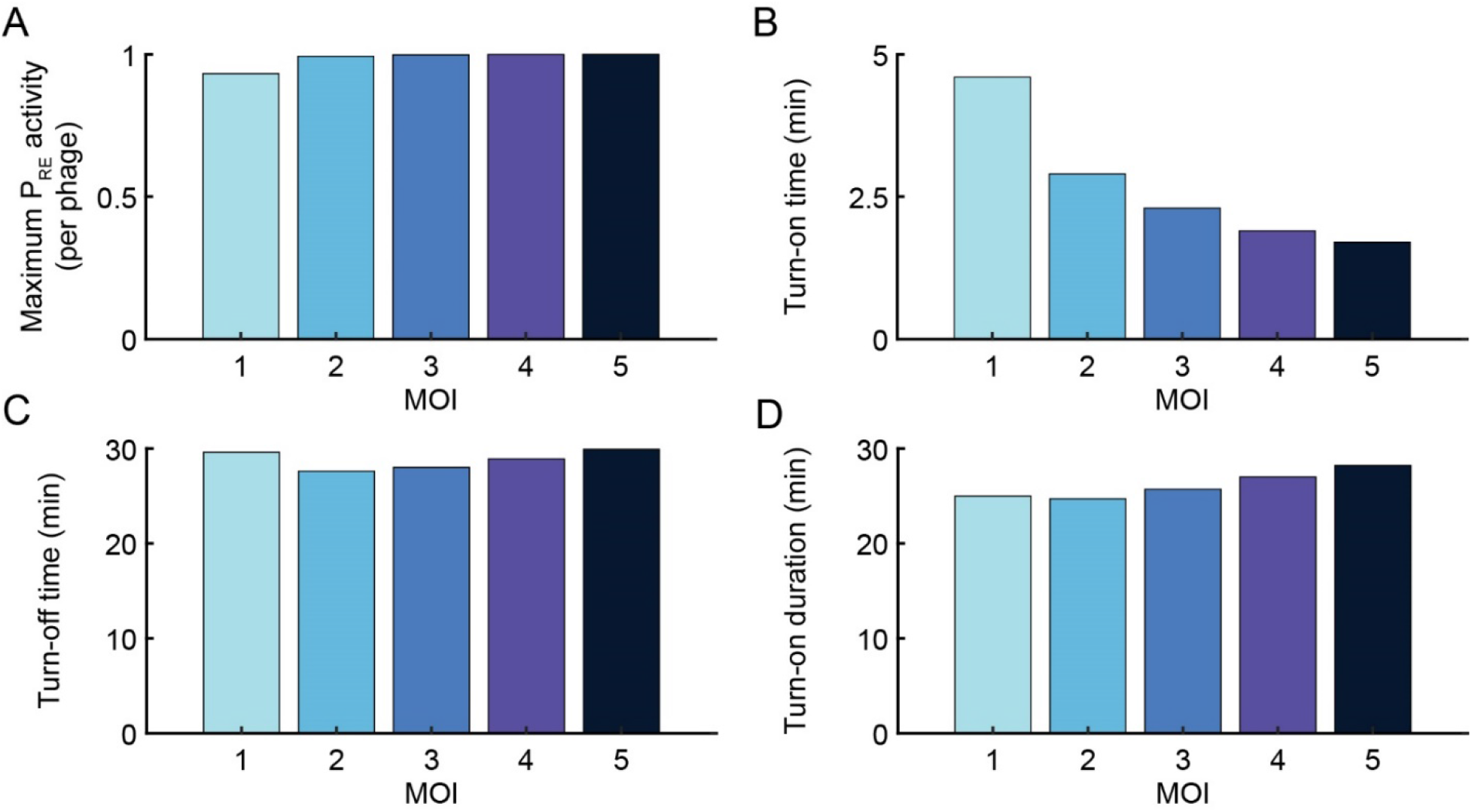
Single phage P_RE_ activity during *P*+ infection depends only weakly on MOI. Model-predicted CII-activated *cI* transcription from P_RE_, at the single phage level, shows weak MOI-dependence of its **(A)** amplitude, **(B)** turn-on time, **(C)**, turn-off time (τ_PRE_), and **(D)** turn-on duration. The turn-on (turn-off) time is defined as the first (last) time that P_RE_ activity is greater than or equal to 10% of its maximum value, while turn-on duration is the difference between these times.

**Figure S11.**
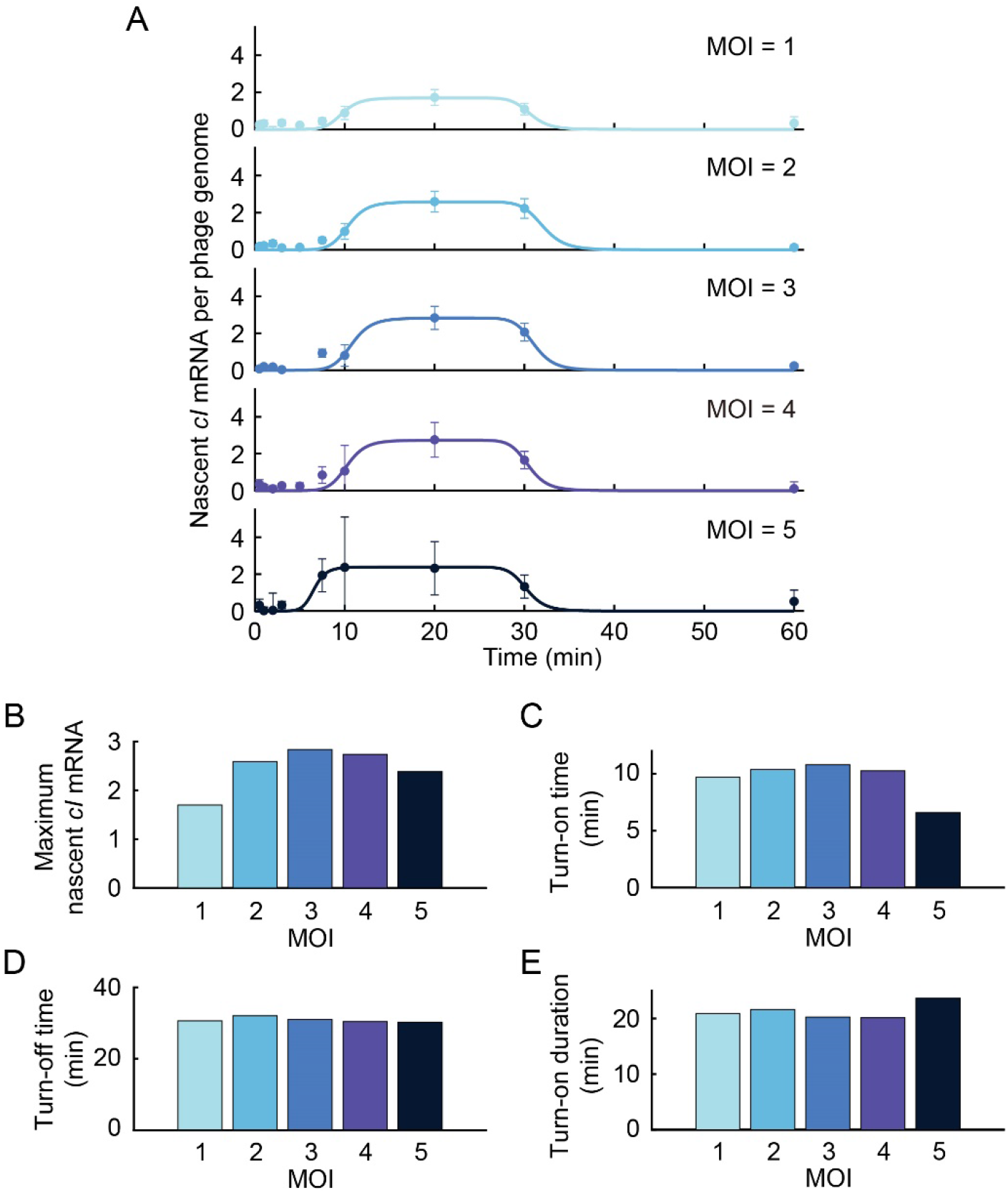
Measured kinetics of nascent *cI* mRNA. **(A)** The number of nascent *cI* mRNA per phage genome following infection by λ *cI*857 *Pam*80 *PlparS*, at single-cell MOI of 1–5. The infection procedure is described in **Experimental Methods Section 4.2**, the mRNA quantification and single-cell MOI identification in **Experimental Methods Sections 8.1** and **8.2**, respectively. Nascent mRNA was quantified based on colocalization of the smFISH and ParB signals, following the method of Wang, M., et al., Nat Microbiol, 2019. The turn-on and turn-off of *cI* transcription were both fitted to a Hill function with coefficient *h* = 10 (solid lines). **(B)** The maximum number of nascent *cI* mRNA per phage genome as a function of MOI. **(C-D)** The time of turn-on and of turn-off of *cI* transcription, estimated using the midpoint of the fitted Hill curves, as a function of MOI. **(E)** The duration of *cI* transcription pulse, estimated using the time interval between activation and repression time, as a function of MOI.

**Figure S12.**
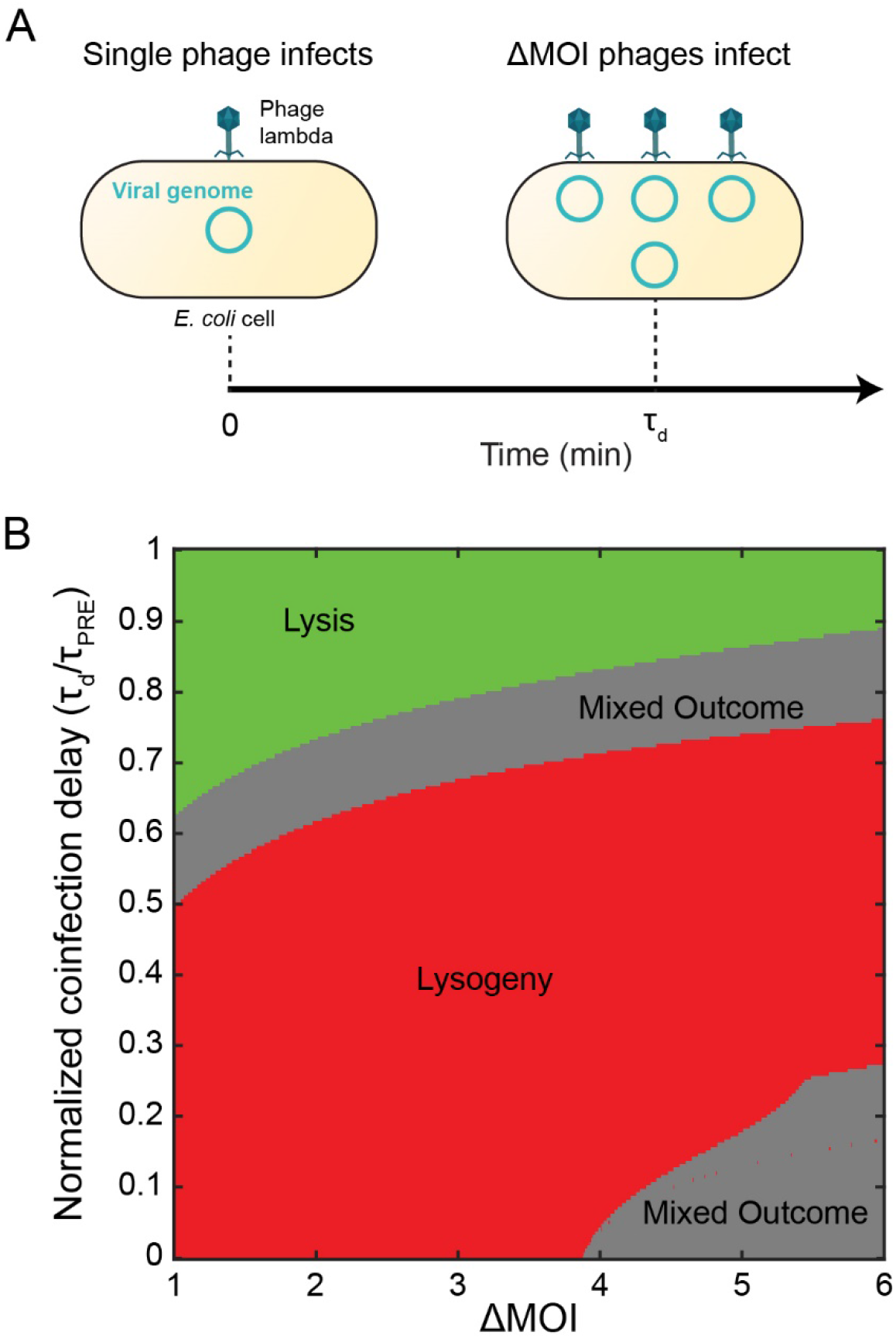
Coinfection delays of τ_pre_, still result in lysis even when the number of coinfecting viruses is greater than 2. **(A)** Following infection with a single phage, ΔMOI additional phages infect at t = Td. **(B)** Model-predicted infection outcome as a function of coinfection delay Td and ΔMOI following MOI = 1 infection by *P*+ phage. Even when ΔMOI is 3-fold larger than MOI*=2, the critical MOI at which the system transitions to lysogeny during simultaneous coinfection (**Figures 3B** and **6D**, main text), only coinfection delays below τ_pre_, (the time when P_RE_ activity falls below 10% of its maximum possible value; see main text) result in lysogeny.

**Figure S13.**
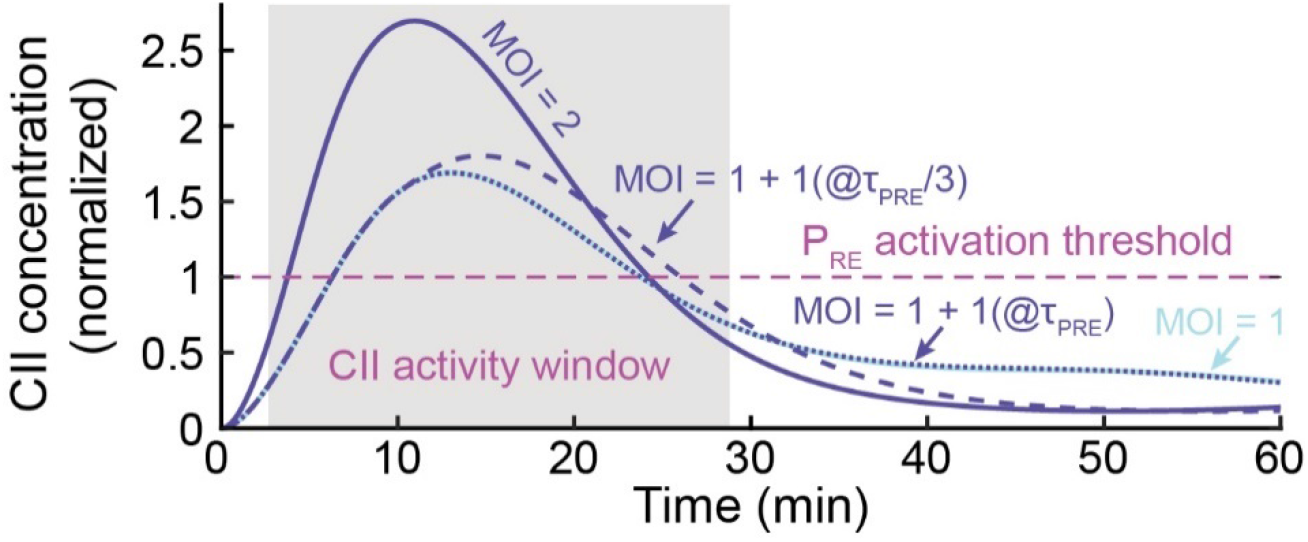
Delayed infection does not result in a second pulse of P_RE_ activity. Model-predicted CII concentration during infection by *P*+ phages for 4 scenarios: Infection by a single phage (light blue solid line), simultaneous infection by two phages (dark blue solid line), infection by a single phage followed by a second phage at time τ_pre_/3 (dark blue dashed line), and infection by a single phage followed by a second phage at time τ_pre_, (dark blue dotted line). The addition of a second phage after a delay (dark blue dashed and dotted lines) does not result in a second pulse of CII. CII concentration is normalized by the P_RE_ activation threshold, and the MOI-averaged CII activity window (defined as the time span during which P_RE_ activity is at least 10% of its maximum possible value) is indicated by the gray shading.

**Figure S14.**
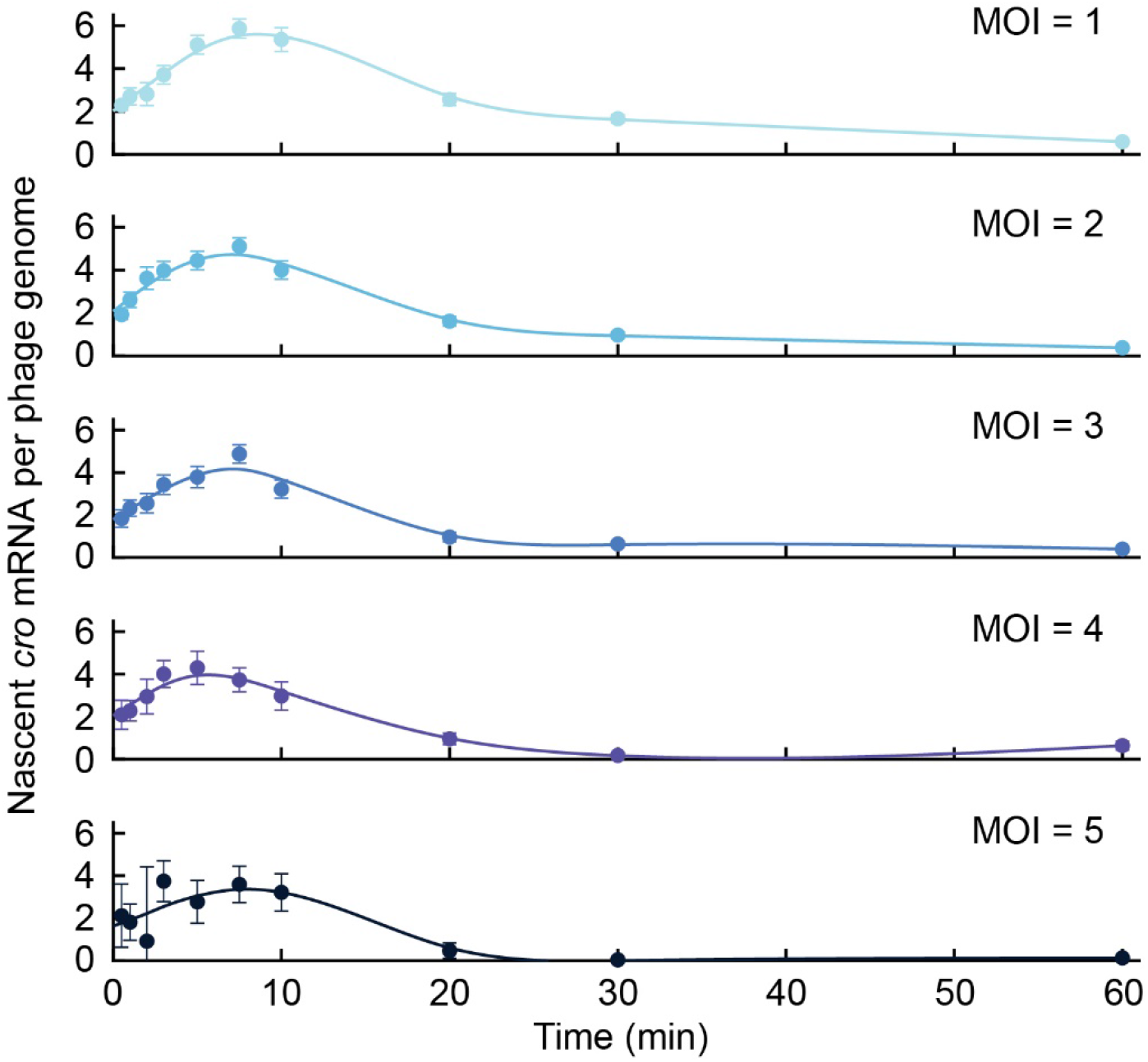
Measured kinetics of nascent *cro* mRNA. The number of nascent *cro* mRNA per phage genome following infection by λ *cI*857 *Pam*80 *PlparS*, at single-cell MOI of 1-5. The infection procedure is described in **Experimental Methods Section 4.2**, the mRNA quantification and single-cell MOI identification in **Experimental Methods Sections 8.1** and **8.2**, respectively. Nascent mRNA was quantified based on colocalization of the smFISH and ParB signals, following the methods of Wang, M., et al., Nat Microbiol, 2019. Solid lines are splines, used to guide the eye.

**Figure S15.**
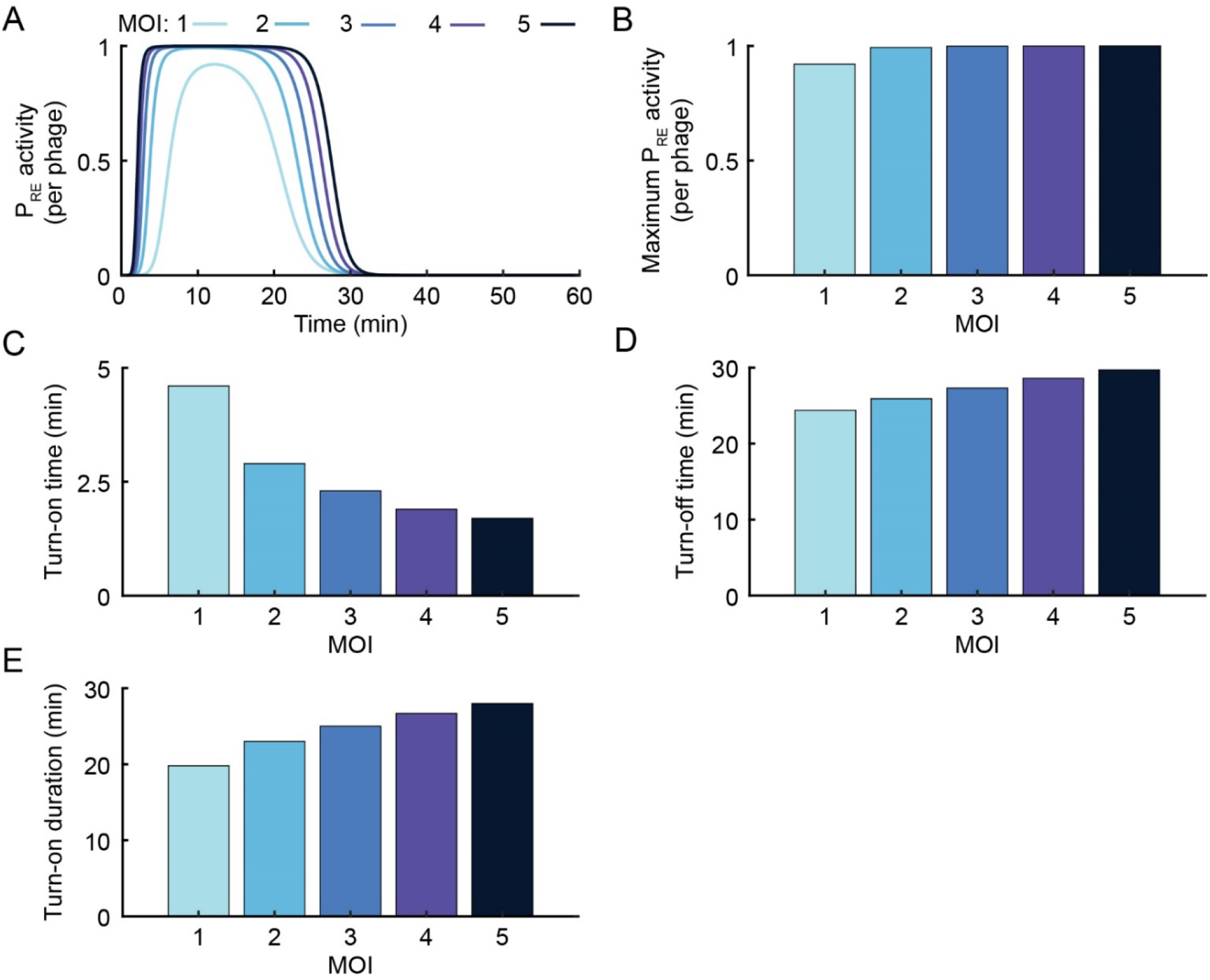
CII activation of P_RE_ during infection by *P*- phage is only weakly MOI-dependent. **(A)** Model-predicted activity of the P_RE_ promoter following infection by *P*- phage at MOI = 1–5. Similar to P_RE_ activity following infection by *P*+ phage (**Figure TS5**; also see **Figure 4A**, main text), P_RE_ activity during *P*- infection does not show strong MOI-dependence in **(B)** amplitude, **(C)** turn-on time, **(D)**, turn-off time (τ_pre_), or **(E)** turn-on duration. The turn-on (turn-off) time is defined as the first (last) time that P_RE_ activity is greater than or equal to 10% of its maximum value, while turn-on duration is the difference between these times.

**Figure S16.**
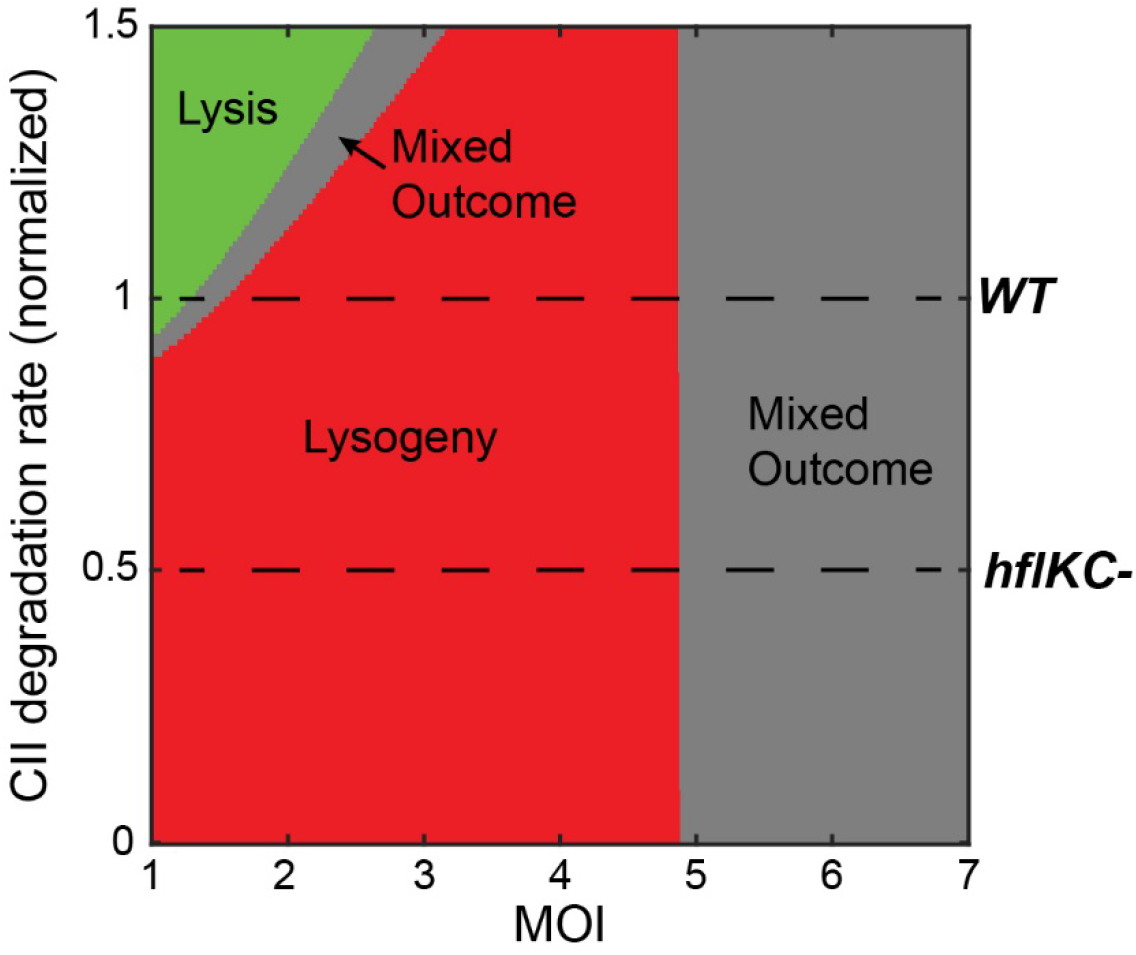
Modulation of CII’s degradation rate can generate lysogenic outcomes at MOI = 1. Model-predicted infection outcome as a function of MOI and CII degradation rate (normalized by the fitted wild-type (*hflKC*+) degradation rate). Perturbations which sufficiently decrease the CII degradation rate (*hflKC*- mutants, black dashed line) result in lysogeny even at MOI = 1.

## Supplementary Tables

**Table S1.** Bacterial strains used in this study.

| Strain name | Relevant genotype or description | Source |
| --- | --- | --- |
| MG1655 | Wild-type | Lab stock |
| LE392 | *glnV* (supE44), *tryT* (supF58) | Lab stock |
| TY132 | MG1655 $\Delta hflK$ | Lab stock |
| TY134 | MG1655 $\Delta hflC$ | Lab stock |

**Table S2.**
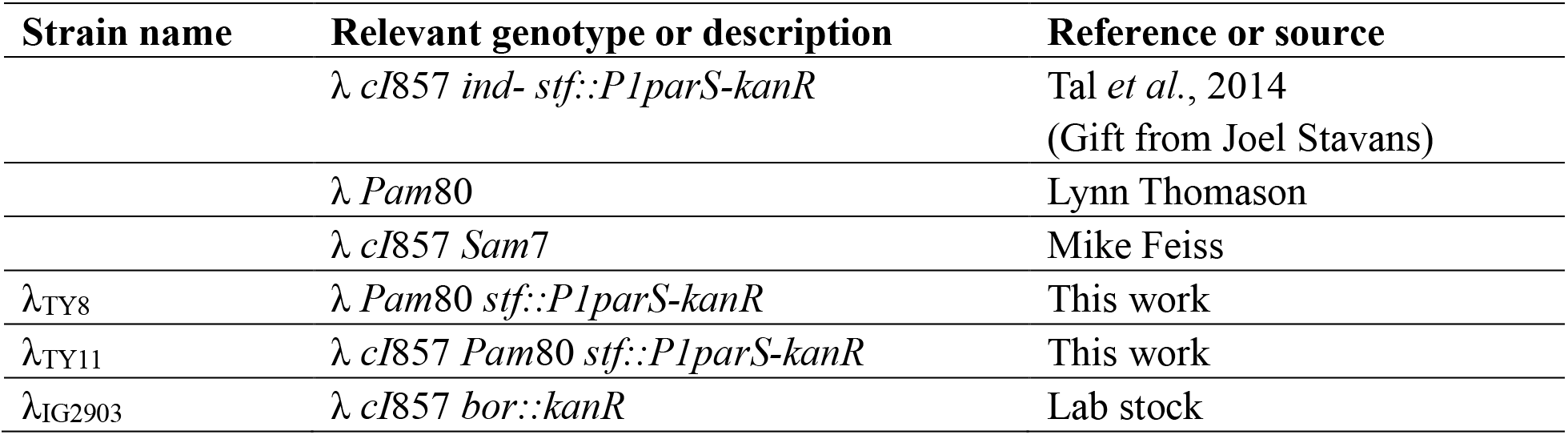
Phage strains used in this study.

**Table S3.** Plasmids used in this study.

| Plasmid name | Description | Reference or source |
| --- | --- | --- |
| pALA3047 | P<sub>lac</sub>-*cfp*-P1-Δ30*parB-ampR* | Stuart Austin |
| pTY001 | Carrying *P1parS-kanR* flanked by<br>lambda *stf* homology | This work |
| pKM208 | P<sub>lac</sub>-*gam-beta-exo-ampR* | Murphy and Campellone, 2003 |

**Table S4.**
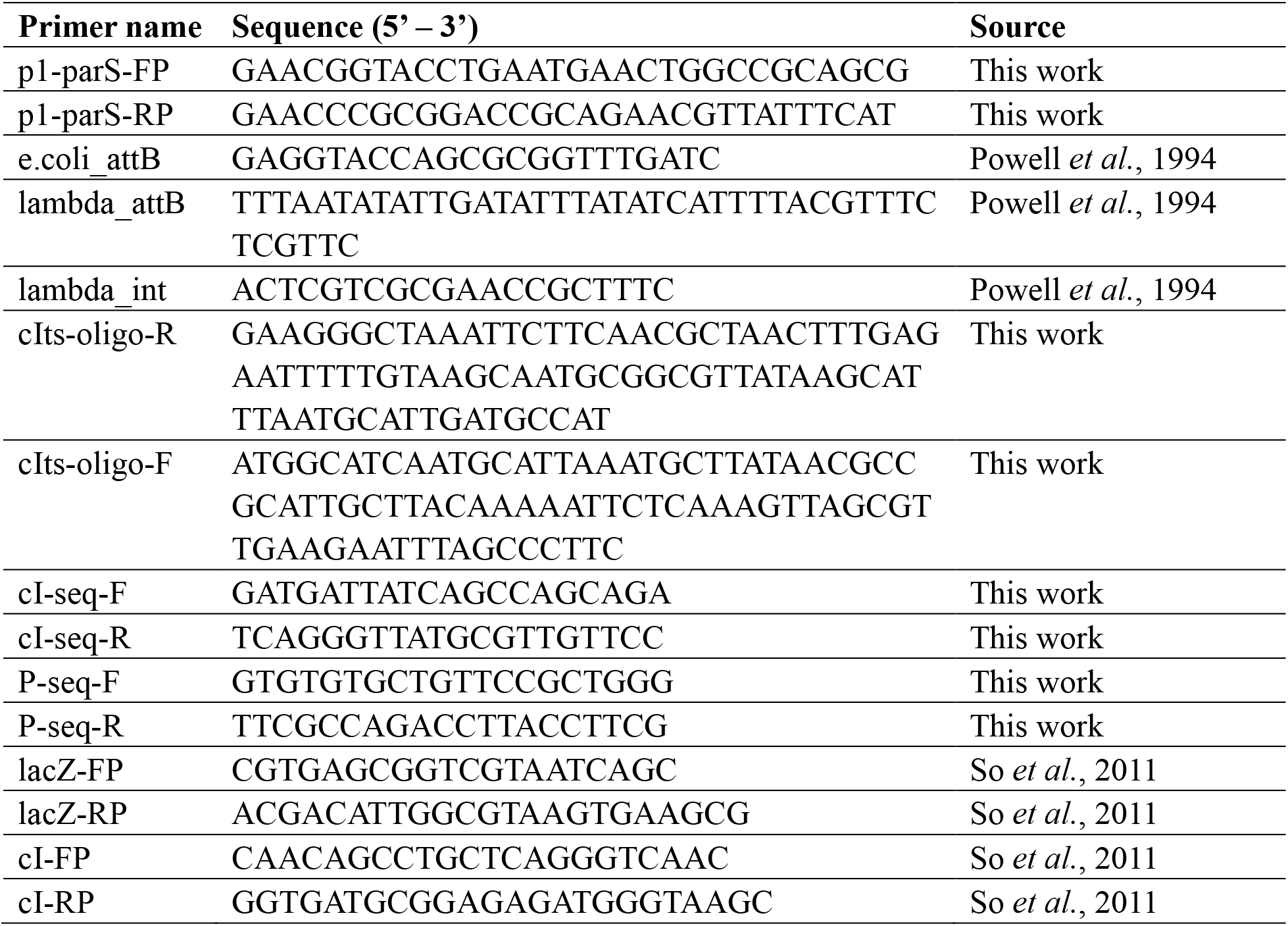
Primers used in this study.

**Table S5.**
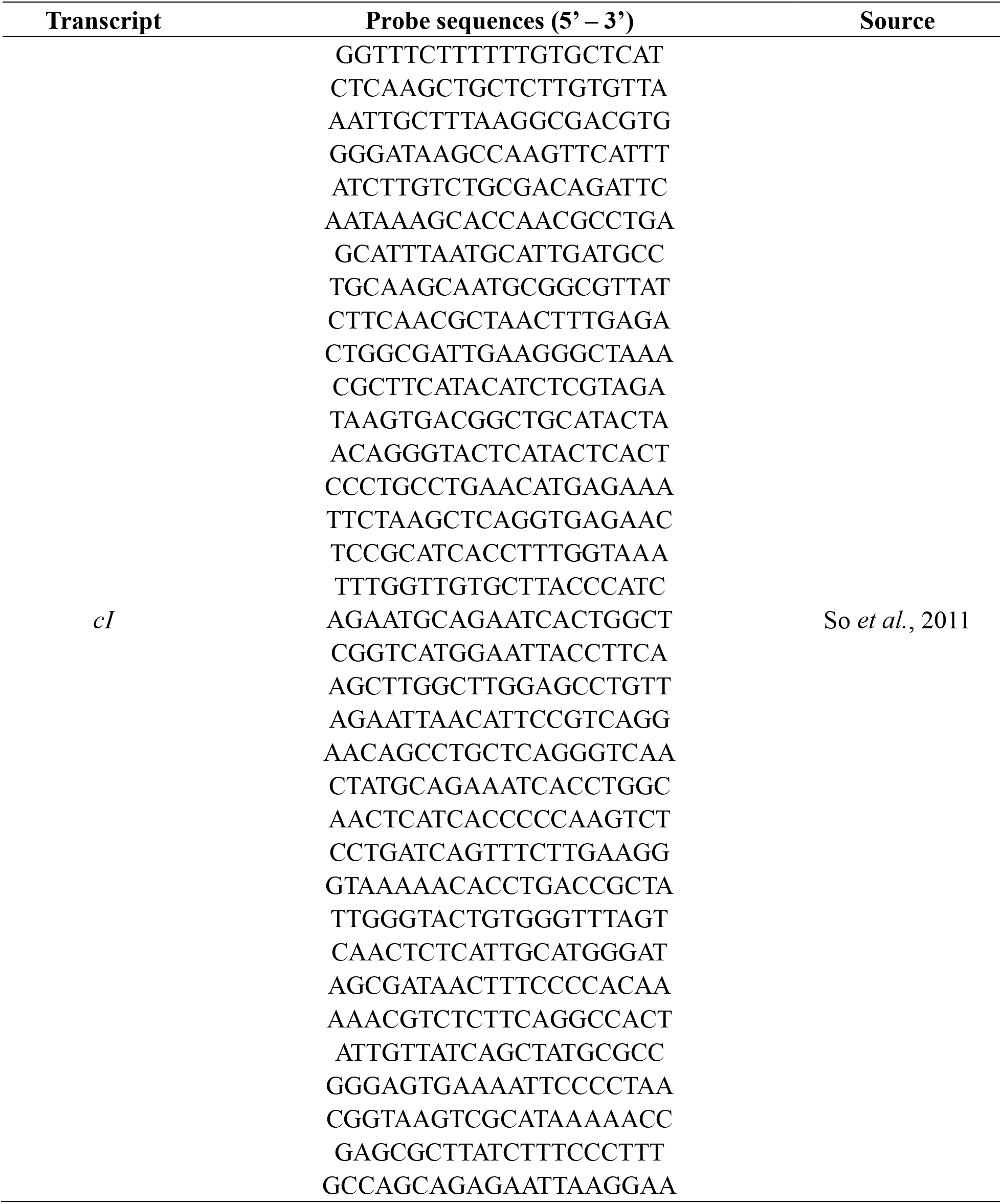
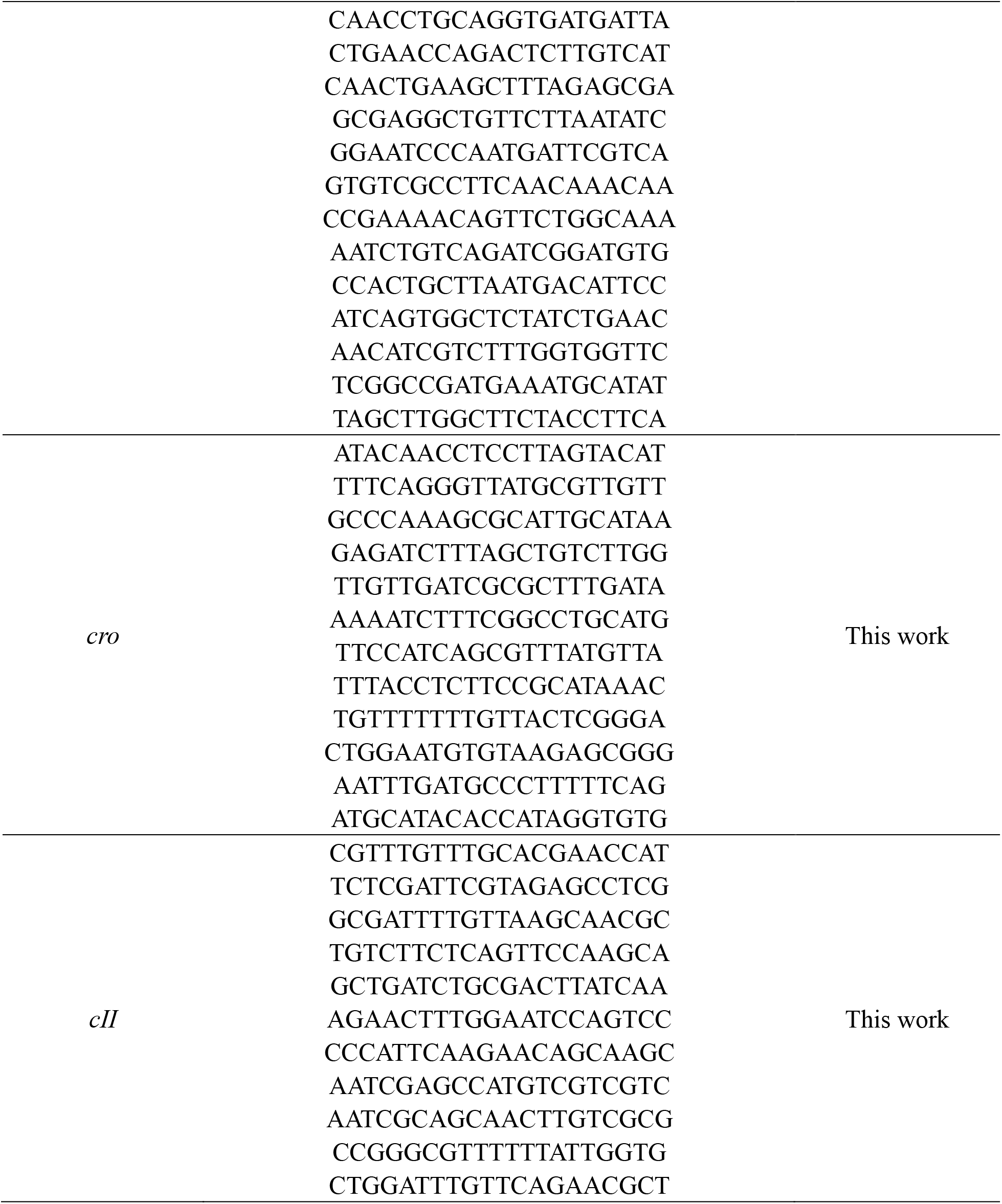
DNA oligos used for smFISH.

**Table S6.** Sample sizes for single-cell infection experiments.

| **Dataset 1** |  |  |  |  |  |  |
| --- | --- | --- | --- | --- | --- | --- |
| Time after infection | Total number of cells (including uninfected) | Number of cells with |  |  |  |  |
|  |  | MOI = 1 | MOI = 2 | MOI = 3 | MOI = 4 | MOI = 5 |
| 0.5 min | 452 | 103 | 117 | 76 | 31 | 7 |
| 1 min | 737 | 154 | 159 | 145 | 79 | 42 |
| 2 min | 355 | 100 | 85 | 56 | 33 | 12 |
| 3 min | 462 | 104 | 105 | 72 | 57 | 36 |
| 5 min | 351 | 103 | 95 | 36 | 24 | 10 |
| 7.5 min | 219 | 45 | 40 | 43 | 25 | 10 |
| 10 min | 56 | 11 | 8 | 18 | 6 | 1 |
| 20 min | 76 | 28 | 14 | 11 | 1 | 1 |
| 30 min | 94 | 24 | 10 | 12 | 5 | 4 |
| 60 min | 822 | 205 | 186 | 86 | 61 | 43 |

| **Dataset 2** |  |  |  |  |  |  |
| --- | --- | --- | --- | --- | --- | --- |
| Time after infection | Total number of cells (including uninfected) | Number of cells with |  |  |  |  |
|  |  | MOI = 1 | MOI = 2 | MOI = 3 | MOI = 4 | MOI = 5 |
| 0.5 min | 804 | 267 | 159 | 50 | 17 | 3 |
| 1 min | 488 | 99 | 109 | 95 | 43 | 26 |
| 2 min | 299 | 93 | 78 | 42 | 19 | 8 |
| 3 min | 364 | 72 | 66 | 69 | 41 | 31 |
| 5 min | 664 | 194 | 151 | 89 | 37 | 17 |
| 7.5 min | 832 | 221 | 195 | 122 | 55 | 18 |
| 10 min | 467 | 123 | 108 | 73 | 35 | 21 |
| 20 min | 875 | 214 | 188 | 120 | 91 | 45 |
| 30 min | 1387 | 338 | 223 | 174 | 142 | 94 |
| 60 min | 820 | 202 | 167 | 120 | 65 | 32 |

